# Charting extracellular transcriptomes in The Human Biofluid RNA Atlas

**DOI:** 10.1101/823369

**Authors:** Eva Hulstaert, Annelien Morlion, Francisco Avila Cobos, Kimberly Verniers, Justine Nuytens, Eveline Vanden Eynde, Nurten Yigit, Jasper Anckaert, Anja Geerts, Pieter Hindryckx, Peggy Jacques, Guy Brusselle, Ken R. Bracke, Tania Maes, Thomas Malfait, Thierry Derveaux, Virginie Ninclaus, Caroline Van Cauwenbergh, Kristien Roelens, Ellen Roets, Dimitri Hemelsoet, Kelly Tilleman, Lieve Brochez, Scott Kuersten, Lukas Simon, Sebastian Karg, Alexandra Kautzky-Willers, Michael Leutner, Christa Nöhammer, Ondrej Slaby, Roméo Willinge Prins, Jan Koster, Steve Lefever, Gary P. Schroth, Jo Vandesompele, Pieter Mestdagh

## Abstract

Extracellular RNAs present in biofluids have emerged as potential biomarkers for disease. Where most studies focus on plasma or serum, other biofluids may contain more informative RNA molecules, depending on the type of disease. Here, we present an unprecedented atlas of messenger, circular and small RNA transcriptomes of a comprehensive collection of 20 different human biofluids. By means of synthetic spike-in controls, we compared RNA content across biofluids, revealing a more than 10 000-fold difference in RNA concentration. The circular RNA fraction is increased in nearly all biofluids compared to tissues. Each biofluid transcriptome is enriched for RNA molecules derived from specific tissues and cell types. In addition, a subset of biofluids, including stool, sweat, saliva and sputum, contains high levels of bacterial RNAs. Our atlas enables a more informed selection of the most relevant biofluid to monitor particular diseases. To verify the biomarker potential in these biofluids, four validation cohorts representing a broad spectrum of diseases were profiled, revealing numerous differential RNAs between case and control subjects. Taken together, our results reveal novel insights in the RNA content of human biofluids and may serve as a valuable resource for future biomarker studies. All spike-normalized data is publicly available in the R2 web portal and serve as a basis to further explore the RNA content in biofluids.

## Introduction

Extracellular RNAs (exRNAs) in blood and other biofluids are emerging as potential biomarkers for a wide range of diseases^1–6^. These so-called liquid biopsies may offer a non-invasive alternative to tissue biopsies for both diagnosis and treatment response monitoring. Previous studies have extensively profiled the small RNA content of several biofluids and identified large differences in the small RNA content amongst different biofluids.^1–12^ These efforts were gathered by the NIH Extracellular RNA Communication Consortium in the exRNA Atlas Resource (https://exrna-atlas.org).^8^ Besides microRNAs (miRNAs), the most studied small RNA biotype in biofluids, other small RNAs, such as piwi‐interacting RNAs (piRNAs), small nuclear RNAs (snRNAs), small nucleolar RNAs (snoRNAs), ribosomal RNAs (rRNAs), transfer RNA fragments (tRNAs) and Y‐RNAs have also been identified^5–7,9,12,13^. Weber et al.^13^ was the first to compare the miRNA content in 12 different human biofluids (pooled samples of plasma, saliva, tears, urine, amniotic fluid, colostrum, breast milk, bronchial lavage fluid, cerebrospinal fluid, peritoneal fluid, pleural fluid and seminal plasma) using reverse transcription quantitative polymerase chain reaction (RT-qPCR) of selected miRNAs. Large variations in RNA concentration were observed among the different biofluids, with the highest small RNA concentrations measured in breast milk and seminal fluid. Since the advent of small RNA sequencing, other small RNA biotypes were characterized in various biofluids, such as plasma, serum, stool, urine, amniotic fluid, bronchial lavage fluid, bile, cerebrospinal fluid (CSF), saliva, seminal plasma and ovarian follicle fluid^5,7,9,9,12^. The distribution of small RNA biotypes clearly varies across these biofluids, with a high abundance of piRNAs and tRNAs reported in urine and a high abundance of Y-RNAs in plasma^6,7,12^. Also non-human RNA sequences, mapping to bacterial genomes, were reported in plasma, urine and saliva^6^.

A systematic RNA-sequencing analysis of biofluids to explore the messenger RNAs (mRNA) and circular RNA (circRNA) transcriptome is challenging due to low RNA concentration and RNA fragmentation in biofluids. As such, most studies have explored the abundance of individual mRNAs in one specific biofluid by RT-qPCR^14–20^. CircRNAs have been reported in saliva^21^, semen^22^, blood^23^ and urine^24,25^. Recently, the mRNA content of plasma and serum has been investigated using dedicated sequencing approaches like Phospho-RNA-Seq, SILVER-seq and SMARTer Stranded Total RNA-Seq method^26–29^. Studies comparing the small RNA, mRNA and circRNA content in a wide range of human biofluids are currently lacking and are essential to explore the biomarker potential of exRNAs.

The goal of the Human Biofluid RNA Atlas is to define the extracellular transcriptome across a wide range of human biofluids (amniotic fluid, aqueous humor, ascites, bile, bronchial lavage fluid, breast milk, cerebrospinal fluid, colostrum, gastric fluid, pancreatic cyst fluid, plasma, saliva, seminal fluid, serum, sputum, stool, synovial fluid, sweat, tear fluid and urine) and to assess biomarker potential in selected case-control cohorts. We used small RNA-sequencing to quantify different small RNA species and present a dedicated mRNA-capture sequencing workflow to simultaneously quantify mRNAs and circRNAs.

In the first phase of our study, small RNA sequencing and mRNA capture sequencing was performed in a discovery cohort of 20 different biofluids (Fig. 1). The goal of this phase was to assess the technical feasibility of the methodology and to generate a comprehensive set of mRNAs, circRNAs and small RNAs in which the contributing tissues and cell types per biofluid were assessed.

**Fig. 1.**
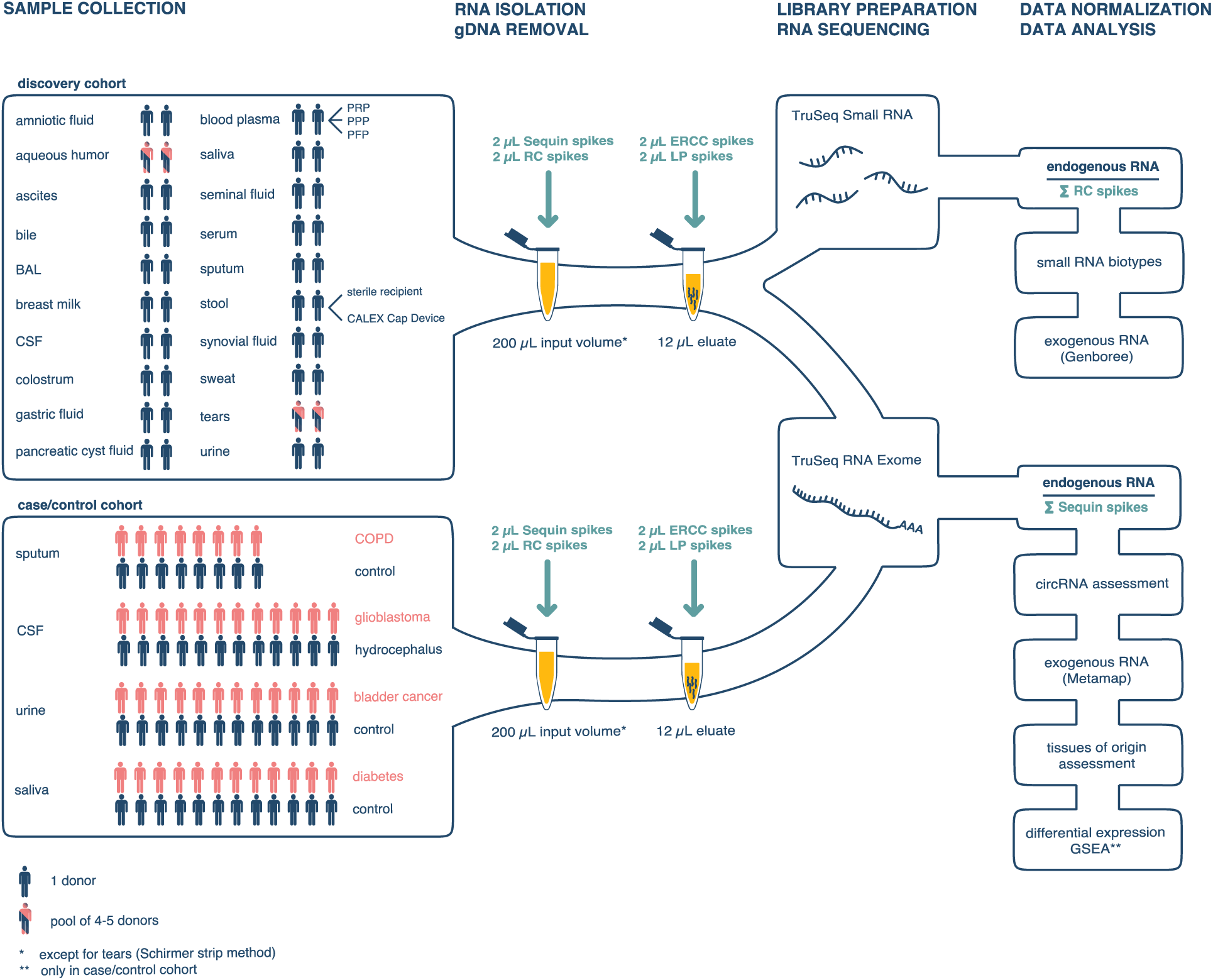
Study flow chart. In the discovery cohort, 20 different biofluids were collected in two donors or in a pool of 4-5 donors. In the case/control cohorts, selected biofluids (sputum, CSF, urine and saliva) were collected in 8-12 patients and an equal number of healthy controls. Both small RNA sequencing and mRNA capture sequencing were performed in the discovery cohort. In the case/control cohorts, mRNA capture sequencing was performed. To compare the RNA content across the different biofluids, the RC spikes and the Sequin spikes are used for normalization of small RNA and mRNA data, respectively. BAL, bronchoalveolar lavage fluid; CSF, cerebrospinal fluid; PRP, platelet-rich plasma; PPP, platelet-poor plasma; PFP, platelet-free plasma

In the second phase of our study, we aimed to investigate the biological relevance of exRNAs in various biofluids. Therefore, mRNA capture sequencing was applied to four different case/control cohorts, each consisting of 16-24 samples (Fig. 1). These samples included sputum samples from 8 patients with chronic obstructive pulmonary disease (COPD) versus 8 controls, urine samples from 12 bladder cancer patients versus 12 controls, CSF samples from 12 glioblastoma patients versus 12 hydrocephalus patients and saliva samples from 12 diabetes mellitus patients versus 12 controls.

The resulting catalog of extracellular transcriptomes of 185 human samples can guide researchers in the biomarker field to investigate other biofluids besides the well-studied blood-derived ones and is a first step to more dedicated mRNA and circRNA profiling of biofluids in larger cohorts.

## Results

### RNA spike-in controls enable process control of the RNA sequencing workflow

Synthetic spike-in RNA sequences are crucial to control the process from RNA isolation to RNA sequencing, especially when working with challenging and low input material. We applied 4 different mixes of synthetic RNA spike-in controls (in total 189 RNAs) as workflow processing and normalization controls that enable direct comparison of the RNA profiles across the different biofluids. Sequin and Small RNA extraction Control (RC) spikes were added prior to RNA isolation whereas External RNA Control Consortium (ERCC) spikes and small RNA Library Prep (LP) spikes were added to the RNA eluate prior to genomic DNA (gDNA) removal (Fig. 1). Of note, every spike mix consists of multiple RNA molecules of different lengths over a wide concentration range. Detailed information is provided in Supplementary Note 1. Besides normalization, the spike-in controls enabled quality control of the RNA extraction and library preparation steps in the workflow and relative quantification of the RNA yield and concentration across the different biofluids.

First, the correlation between the expected and the observed relative quantities for all four spike mixes can be used to assess quantitative linearity. In the discovery cohort, the expected and the observed relative quantities for all four spike mixes were well correlated (Pearson correlation coefficients range from 0.50 to 1.00 for Sequin spikes, 0.92 to 1.00 for ERCC spikes, 0.44 to 0.98 for RC spikes and 0.40 to 0.96 for LP spikes). In some biofluids (e.g. seminal plasma and tears), the sequencing coverage of spikes was low due to a high concentration of endogenous RNA. Detailed information per sample is provided in Supplementary Fig. 1.

The spike-in controls can also be used to assess the RNA isolation efficiency. The Sequin/ERCC ratio and the RC/LP ratio reflect the relative mRNA and microRNA isolation efficiency, respectively. A 170-fold and 104-fold difference in RNA isolation efficiency across the samples was observed when assessing long and small RNAs, respectively (Supplementary Fig. 2). These differences underline the challenges of working with heterogenous samples and the importance of spike-in controls for proper data normalization and cross-sample comparison of results.

Finally, the spikes can be utilized to normalize the endogenous RNA abundance data. In this study, we applied a biofluid volume-based normalization by dividing the RNA reads consumed by the endogenous transcripts by the sum of the Sequin spikes for mRNA data and by the sum of the RC spikes for small RNA data. The spike-normalized data represent relative abundance values of RNA molecules proportional to the input volume. Of note, there is an inverse relationship between the number of spike-in RNA reads and the number of endogenous RNA reads. As such, the ratio between the sum of the reads consumed by the endogenous transcripts and the total number of spike-in reads is a relative measure for the RNA concentration of the various samples.

### Highly variable mRNA and small RNA content among biofluids in the discovery cohort

Both small RNAs and mRNAs were quantified in each of the 20 biofluids in the discovery cohort. Mapping rates varied substantially across the different biofluids (Fig. 2A). In general, the proportion of mapped reads was higher for the mRNA capture sequencing data (further referred to as mRNA data) than for the small RNA sequencing data, in line with the fact that human mRNAs were enriched using biotinylated capture probes during the library preparation. The fraction of mapped reads in the mRNA data ranged from 16% in stool to 97% in seminal plasma. Low mapping rates were observed in stool, in one of the bile samples and in saliva. Mapping rates for samples in the case/control cohorts are in line with these of the discovery cohort (Supplementary Fig. 3A). In the small RNA sequencing data, the proportion of mapped reads ranged from ~7% in stool, saliva and CSF to 95% in platelet-rich plasma (PRP). A 10 000-fold difference in mRNA and small RNA concentration was observed between the lowest concentrated fluids, i.e. platelet-free plasma, urine and CSF, and the highest concentrated biofluids, i.e. tears, seminal plasma and bile (Fig. 2B). The generalizability of the difference in mRNA concentration between highly concentrated biofluids (seminal plasma) and lowly concentrated biofluids (CSF) was confirmed in additional samples (Supplementary Fig. 3B). In the discovery cohort a 5547-fold difference in mRNA concentration is observed between seminal plasma and CSF; in independent validation samples, a similarly large 19 851-fold difference in mRNA concentration is observed between both biofluids. In the discovery cohort, the mRNA and miRNA concentrations were significantly correlated across biofluids (Pearson correlation coefficient 0.76, p-value = 8.5e-10, Fig. 2D). Normalized abundance levels of exRNAs were significantly correlated between biological replicates within each biofluid (Supplementary Fig. 4). The median Pearson correlation coefficient of the mRNA and the small RNA data was 0.84 and 0.92, respectively. While the mRNA and miRNA data was well correlated in most biofluids (e.g. tears, colostrum, saliva), correlation in other biofluids (e.g. bile, pancreatic cyst fluid) was poor. These biofluids are obtained with a more challenging collection method involving echo-endoscopy, impacting the reproducibility of collection and the correlation of the RNA content between biological replicates.

**Fig. 2.**
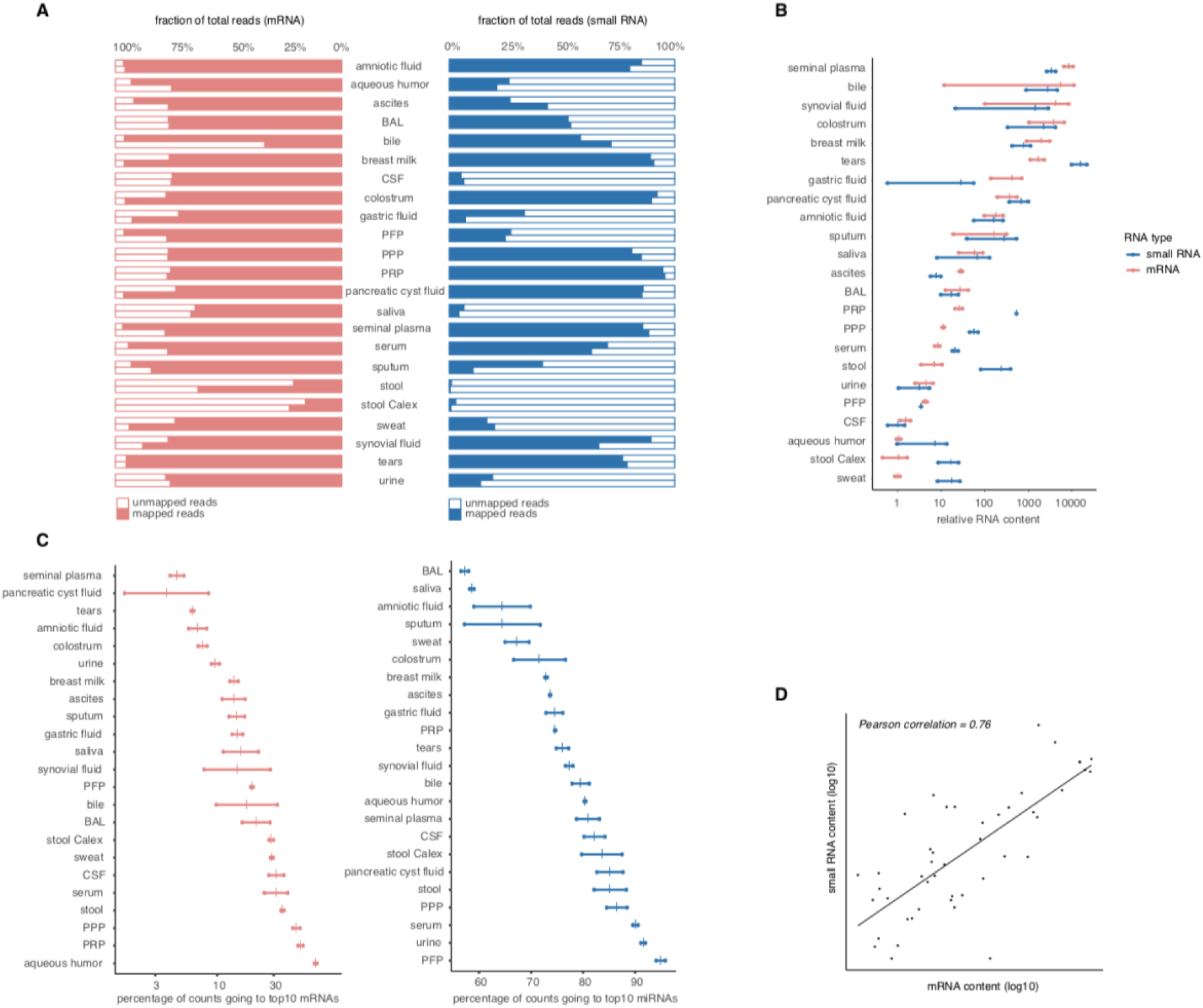
2 mRNA and small RNA content varies across the 20 biofluids. (A) Percentage of the total read count mapping to the human transcriptome. (B) Relative RNA concentration per biofluid; every dot represents the relative RNA concentration in one sample, every vertical mark indicates the mean per biofluid. (C) The diversity of the RNA content expressed as fraction of read counts consumed by the top 10 most abundant mRNAs/miRNAs. Only genes with at least 4 unique reads are taken into account. Every dot represents the fraction in one sample, every vertical mark indicates the mean percentage per biofluid. (D) Correlation between the small RNA and the mRNA relative concentration. The Pearson correlation coefficient is 0.76 (p-value = 8.58 × 10^-10^). The correlation coefficients is calculated on log10 transformed data. BAL, bronchoalveolar lavage fluid; CSF, cerebrospinal fluid; PRP, platelet-rich plasma; PPP, platelet-poor plasma; PFP, platelet-free plasma

The likelihood of identifying RNA biomarkers in a given biofluid will not only depend on its relative RNA concentration, but also on its RNA diversity, here approximated by the fraction of read counts consumed by the top 10 most abundant mRNAs/miRNAs (Fig. 2C). In aqueous humor, the top 10 mRNAs represent up to 70% of all reads, indicating that this fluid does not contain a rich mRNA repertoire. In both PRP and PPP, about 50% of all reads go to the top 10 mRNAs. While amniotic fluid has a median RNA concentration, this fluid seems to contain a diverse mRNA profile, with only 7% of all reads going to the top 10 mRNAs. When looking into the miRNA data, the top 10 miRNAs represent more than 90% of all reads in PFP, urine and serum. BAL contains the most diverse miRNA repertoire, with 57% of all reads going to the top 10 miRNAs. Similar conclusions with respect to biofluid exRNA diversity can be drawn based on the number of miRNAs/mRNAs representing 50% of the counts (Supplementary Fig. 5). RNA diversity is also reflected by the number of detected exRNAs. The total number of mRNAs and miRNAs detected with at least 4 counts in both samples of the same biofluid ranged from 13 722 mRNAs in pancreatic cyst fluid to 107 mRNAs in aqueous humor and from 231 miRNAs in tears to 18 miRNAs in stool (Table 1).

**Table 1.**
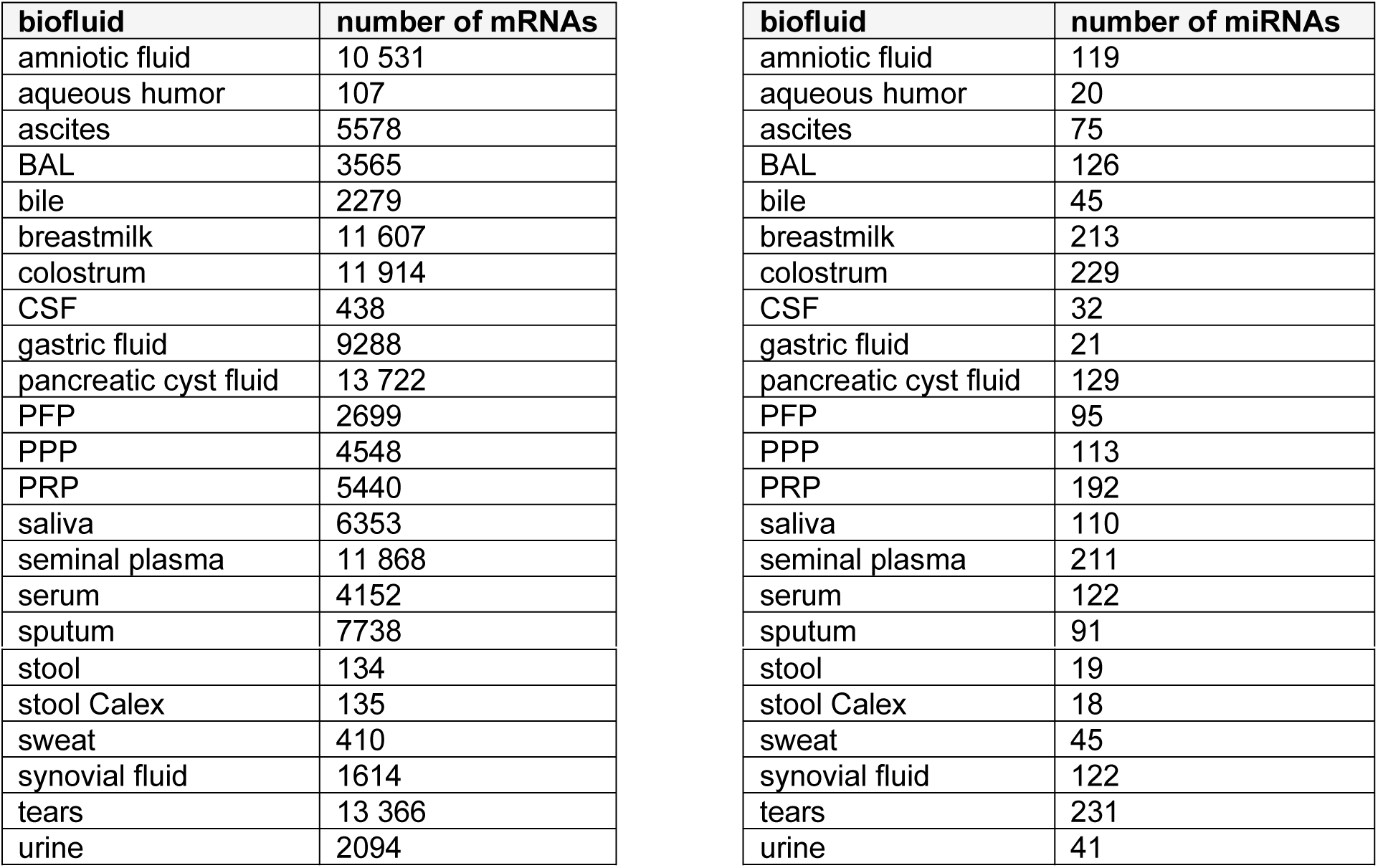
Number of mRNAs and miRNAs per biofluid. The number of mRNAs and miRNAs with at least 4 unique read counts in both replicates is shown per biofluid.

### The distribution of small RNA biotypes varies across the different biofluids

The distribution of small RNA biotypes shows distinct patterns among the 20 different biofluids (Fig. 3). The exceptionally high percentage of miscellaneous RNAs (mainly Y-RNAs) observed in blood-derived fluids is in line with a previous study^12^ and with the Y-RNA function in platelets. The fraction of reads mapping to miRNAs is lower than 15% in all samples but platelet-free plasma and one synovial fluid sample. Tears, bile and amniotic fluid have the highest fraction of tRNA fragments while saliva has the highest fraction of piRNAs. The rRNA fraction is higher than 15% in all samples but tears, aqueous fluid and the three plasma fractions. The majority of these reads map to the 45S ribosomal RNA transcript. The not annotated read fraction contains uniquely mapped reads that could not be classified in one of the small RNA biotypes. These reads most likely originate from degraded longer RNAs, such as mRNAs and long non-coding RNAs.

**Fig. 3.**
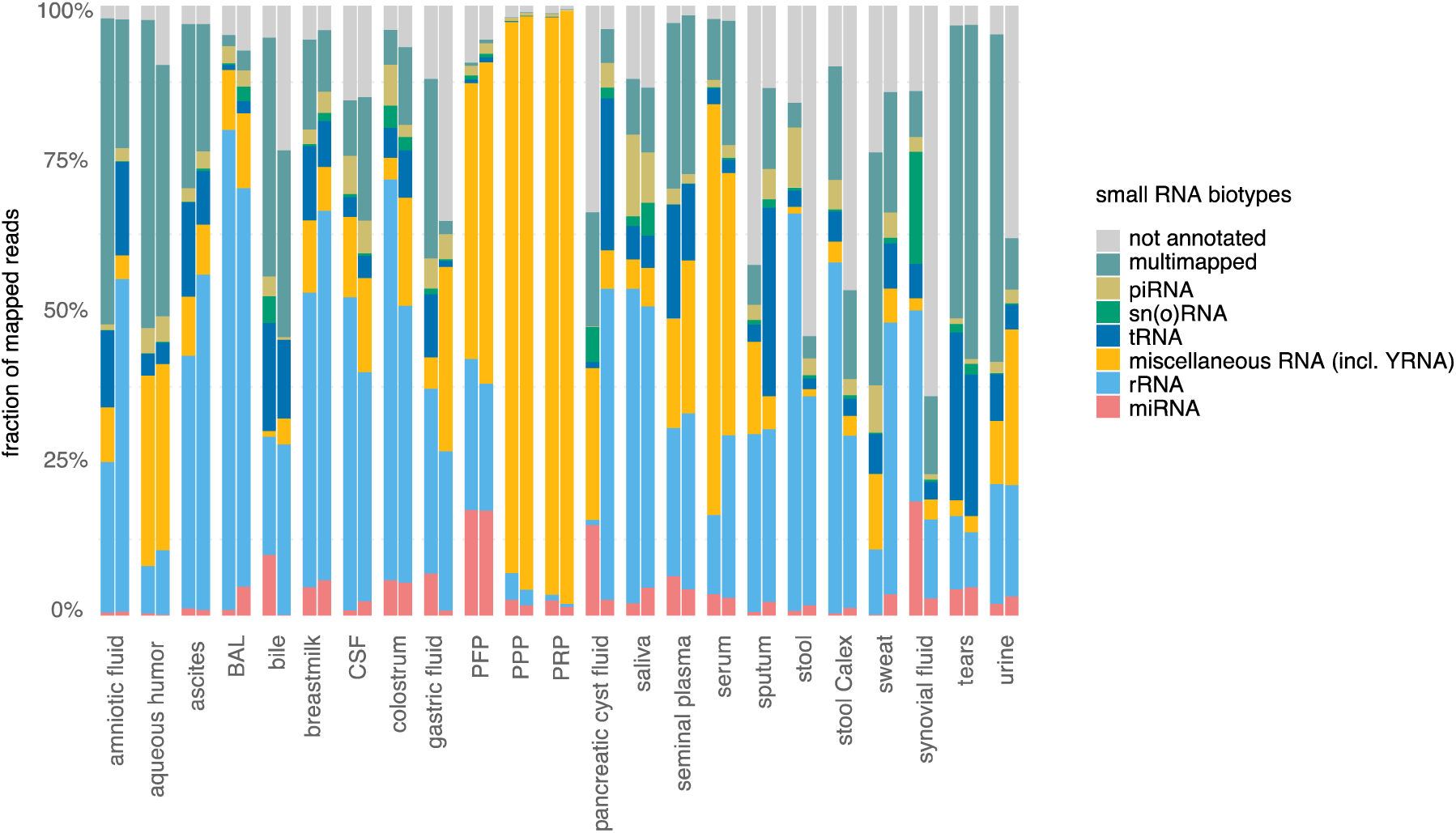
Distinct small RNA biotype patterns are present across the different biofluids. The fraction of reads that align to small RNA biotypes are shown per biofluid. Only mapped reads of the small RNA sequencing data are taken into account. BAL, bronchoalveolar lavage fluid; CSF, cerebrospinal fluid; *miRNA: microRNA;* PFP, platelet-free plasma; PPP, platelet-poor plasma; PRP, platelet-rich plasma; *piRNAs: piwi-interacting RNA; sn(o)RNAs: small nuclear and nucleolar RNAs; tRNAs: transfer RNA*.

### Circular RNAs are enriched in biofluids compared to tissues

CircRNAs are produced from unspliced RNA through a process called back-splicing where a downstream 5’ donor binds to an upstream 3’ acceptor. CircRNAs are resistant to endogenous exonucleases that target free 5’ or 3’ terminal ends. As a result, circRNAs are highly stable and have extended half-lives compared to linear mRNAs.^30^ CircRNAs have been reported to be present in numerous human tissues^24^ and in a few biofluids such as saliva^21^, blood^31^, semen^22^ and urine^24,25^. A direct comparison of the circRNA read fraction between biofluids and tissues is currently lacking in literature. We compared the circRNA fraction, for genes that produce both linear and circular transcripts, identified through mRNA capture sequencing of the 20 biofluids in this study with the circRNA fraction identified in mRNA capture sequencing of 36 cancerous tissue types obtained from the MiOncoCirc Database^24^. While more unique backsplice junctions were identified in tissues compared to biofluids, in line with the higher RNA concentration in tissues (Fig. 4B), the circRNA read fraction is clearly higher in biofluid exRNA compared to cellular RNA (Fig. 4A). The median circRNA read fraction in biofluids is 84.4%, which is significantly higher than the median circRNA read fraction in tissues of 17.5% (Mann-Whitney-U test, two-sided, p-value = 5.36 × 10^-12^). For genes that produce both linear and circular transcripts, the stable circRNAs are more abundant than the linear mRNAs in biofluids, while it is the other way around in tissues.

**Fig. 4.**
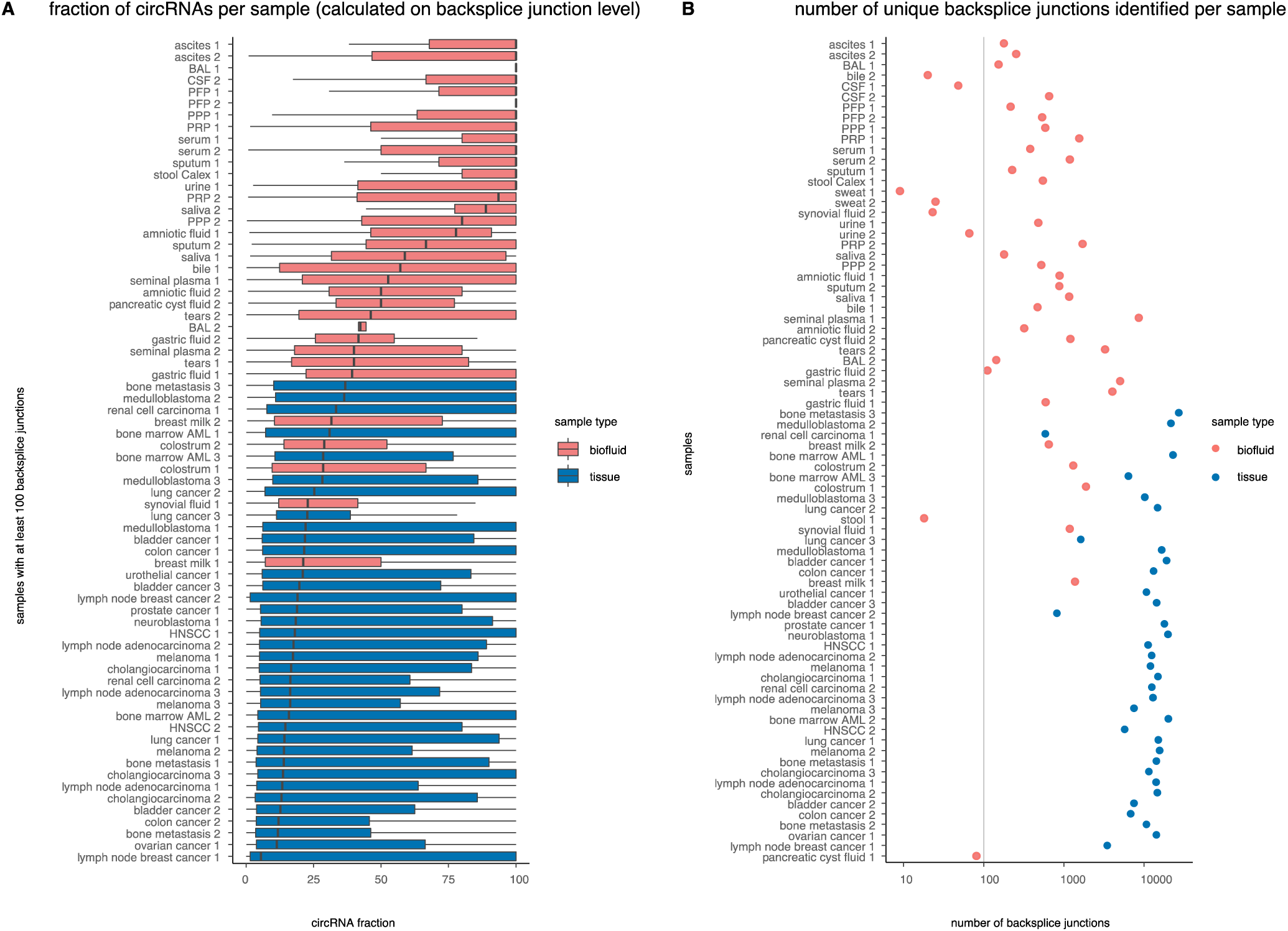
CircRNAs are enriched in biofluids compared to tissues. (A) The circRNA fraction, calculated at the backsplice junction level, is plotted per sample and is higher in cell-free biofluid RNA than in tissue RNA. Only samples with at least 100 backsplice junctions are plotted. (B) The number of unique backsplice junctions per sample is higher in tissues compared to biofluids, in line with the higher input concentration of RNA into the library prep. AML, acute myeloid leukemia; BAL, bronchoalveolar lavage fluid; CSF, cerebrospinal fluid; HNSCC: head and neck squamous-cell carcinoma; PFP, platelet-free plasma; PPP, platelet-poor plasma; PRP, platelet-rich plasma

We used two different methods to define the circRNA read fraction (see *“Circular RNA detection”* in methods; Supplementary Fig. 6): one based on individual backsplice junctions (shown in Fig. 4) and another method based on backsplice junctions aggregated at gene-level (Supplementary Fig. 7). Both methods clearly point towards a substantial enrichment of circRNAs in biofluids.

### Assessment of exogenous RNA in human biofluids

Two dedicated pipelines were used for the non-trivial assessment of the presence of microbial or viral RNA in human biofluid extracellular RNA. Overall, the fraction of bacterial reads is higher in small RNA sequencing data than in the mRNA data, in line with the unbiased nature of small RNA sequencing and the targeted hybrid capture enrichment using probes against human RNA during the mRNA capture library preparation. Stool (both collection methods), sweat, saliva and sputum are among the biofluids with the highest fraction of bacterial RNA in both the small RNA sequencing data and the mRNA data. The percentage of bacterial reads in mRNA data and in small RNA data are significantly correlated across biofluids (Pearson correlation coefficient 0.78, p-value = 1.94e-10).

Bacterial reads in aqueous humor and CSF, two fluids with very low endogenous RNA content that were collected in a sterile setting (and thus presumed to be sterile), most likely reflect background contamination during the workflow^32^. To illustrate the biological relevance of the bacterial signal, we looked into reads mapping to *Campylobacter concisus*, a gram-negative bacterium that is known to primarily colonize the human oral cavity, with some strains translocated to the intestinal tract^33^. We confirm the selective presence of reads mapping to *Campylobacter concisus* in saliva in both the small RNA and the mRNA data(Fig. 5B). In all samples and for both the small RNA and the mRNA data, the percentage of the total reads that maps to viral transcriptomes is less than 1%.

**Fig. 5.**
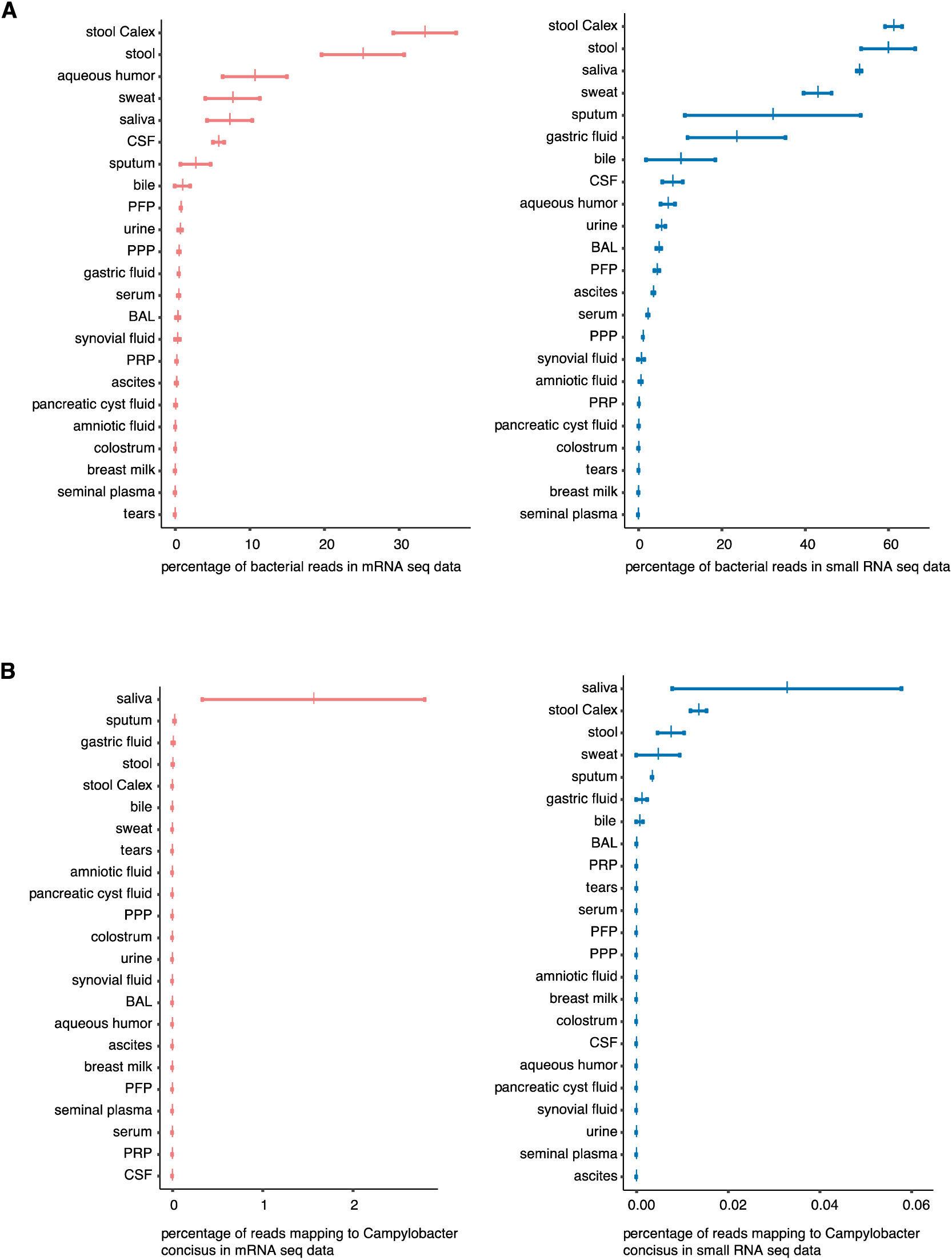
Reads mapping to bacterial genomes. (A) Percentage of reads mapping to bacteria in mRNA data (pink) and in small RNA sequencing data (blue). (B) Percentage of reads mapping to Campylobacter concisus in mRNA data (pink) and in small RNA sequencing data (blue). Campylobacter concisus is known to be present in saliva.

### Assessment of the tissues of origin and deconvolution of pancreatic cyst fluid

Gaining insights in tissue contribution to biofluid RNA profiles may guide the selection of the most appropriate biofluid to investigate a given disease. To define tissues that specifically contribute RNA molecules to individual biofluids, we explored the relationship between extracellular mRNA levels and tissue or cell type specific mRNA signatures. The heatmap in Fig. 6A highlights the relative contribution of tissues and cell types to a specific biofluid compared to the other biofluids. More detailed results per biofluid are shown in Supplementary Fig. 8. The results of this analysis were validated in an independent sample cohort for CSF, saliva, sputum, seminal plasma and urine (Supplementary Fig. 3C). As expected, prostate tissue RNA markers are more abundant in urine and in seminal plasma than in any other biofluid. Both sputum and saliva contain mRNAs specific for trachea and esophagus. In amniotic fluid, markers for esophagus, small intestine, colon and lung are more abundant than the other tissues and cell types, probably reflecting organs that actively shed RNA (at the gestational age of sampling) into the amniotic cavity. These data strongly suggest that biofluid mRNA levels, at least to some degree, reflect intracellular mRNA levels from cells that produce or transport the fluid. To further investigate the origin of biofluid RNA at the cellular level, we applied computational deconvolution of the pancreatic cyst fluid RNA profiles using single cell RNA sequencing data from 10 pancreatic cell types^34^. Fig. 6B reveals that pancreatic cyst fluid 1 consists of 45% of activated stellate cells and 43% of endothelial cells, while pancreatic cyst fluid 2 mainly consists of quiescent stellate cells (38%), endothelial cells (31%) and acinar cells (19%).

**Fig. 6.**
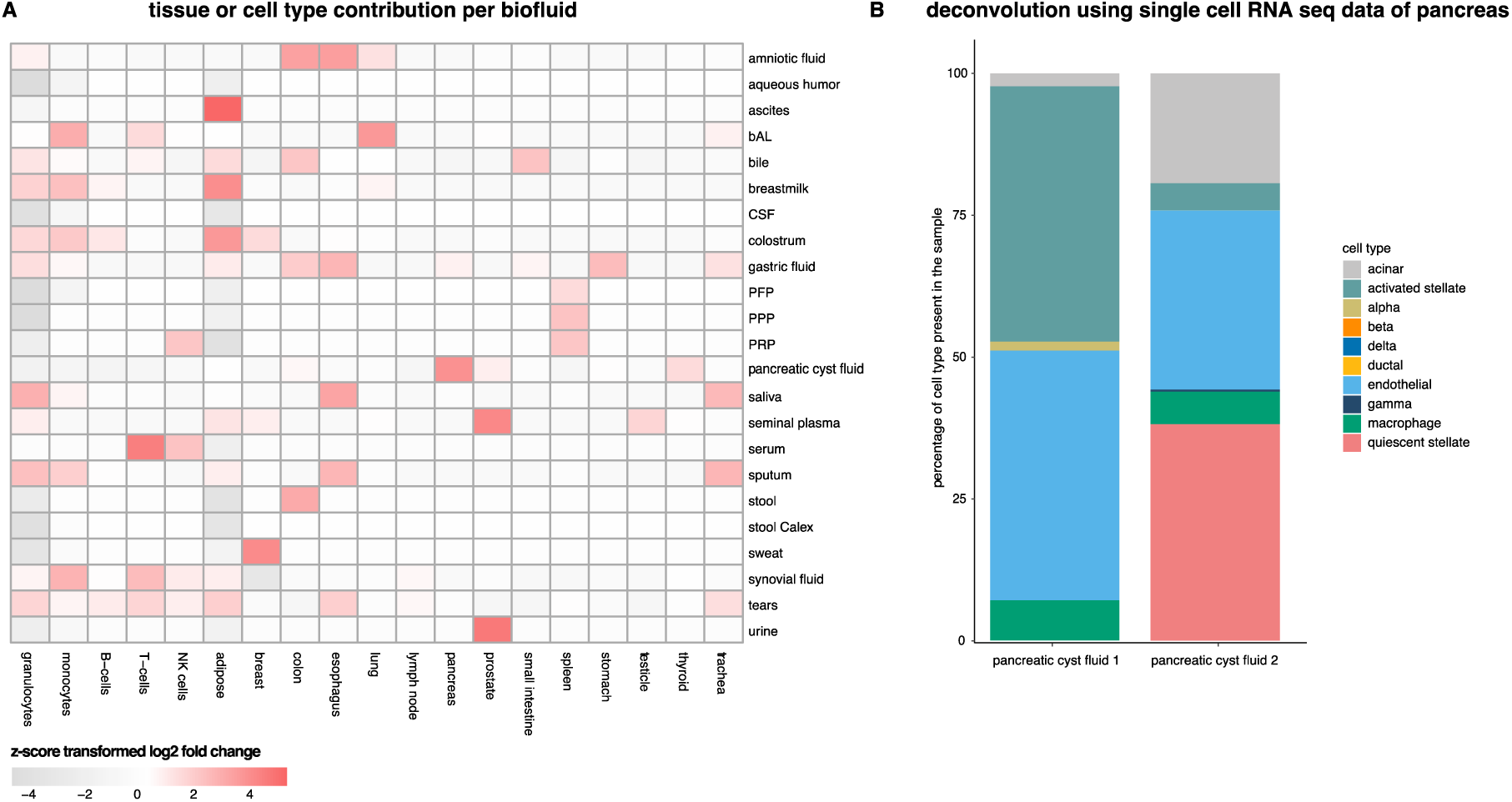
Identification of the tissues of origin per biofluid and deconvolution of pancreatic cyst fluid. (A) Assessment of the tissues of origin in the biofluids of the discovery cohort. Heatmap showing tissues and cell types that contribute more specifically to a certain biofluid compared to the other biofluids. Rows depict the biofluids of the discovery cohort and the columns are the tissues or cell types for which markers were selected based on the RNA Atlas^35^. For visualization purposes, only tissues and cell types with a z-score transformed log2 fold change ≥ |1| in at least one biofluid are shown. (B) Composition of pancreatic cyst fluid samples based on deconvolution using sequencing data from 10 pancreatic cell types.

### Biomarker potential of mRNA in sputum, urine, CSF and saliva in selected case/control cohorts

Additional biofluid samples were collected in patients with a specific disease or in healthy controls to investigate potential biologically relevant differences in mRNA content between both groups. Sequin RNA spikes were used for biofluid volume-based data normalization. Strikingly, the relative RNA concentration in sputum of COPD patients was higher than in non-COPD patients, probably reflecting the high turnover of immune cells during the state of chronic inflammation (Fig. 7A). Differential expression analysis revealed 5513 and 6 mRNAs that were significantly up- and downregulated, respectively, in sputum from COPD patients compared to healthy controls (Fig. 7B). CCL20, the most differential mRNA, showed a 146-fold upregulation in COPD patients compared to healthy donors. This potent chemokine attracting dendritic cells has previously been linked to the pathogenesis of COPD^36,37^. ADA and MMP1, also among the most differential mRNAs, have also been associated with the pathogenesis of COPD^38–40^. To verify the RNA-seq findings, 8/8 of the most differentially abundant mRNAs were validated by RT-qPCR (Supplementary Fig. 9A-B).

**Fig. 7.**
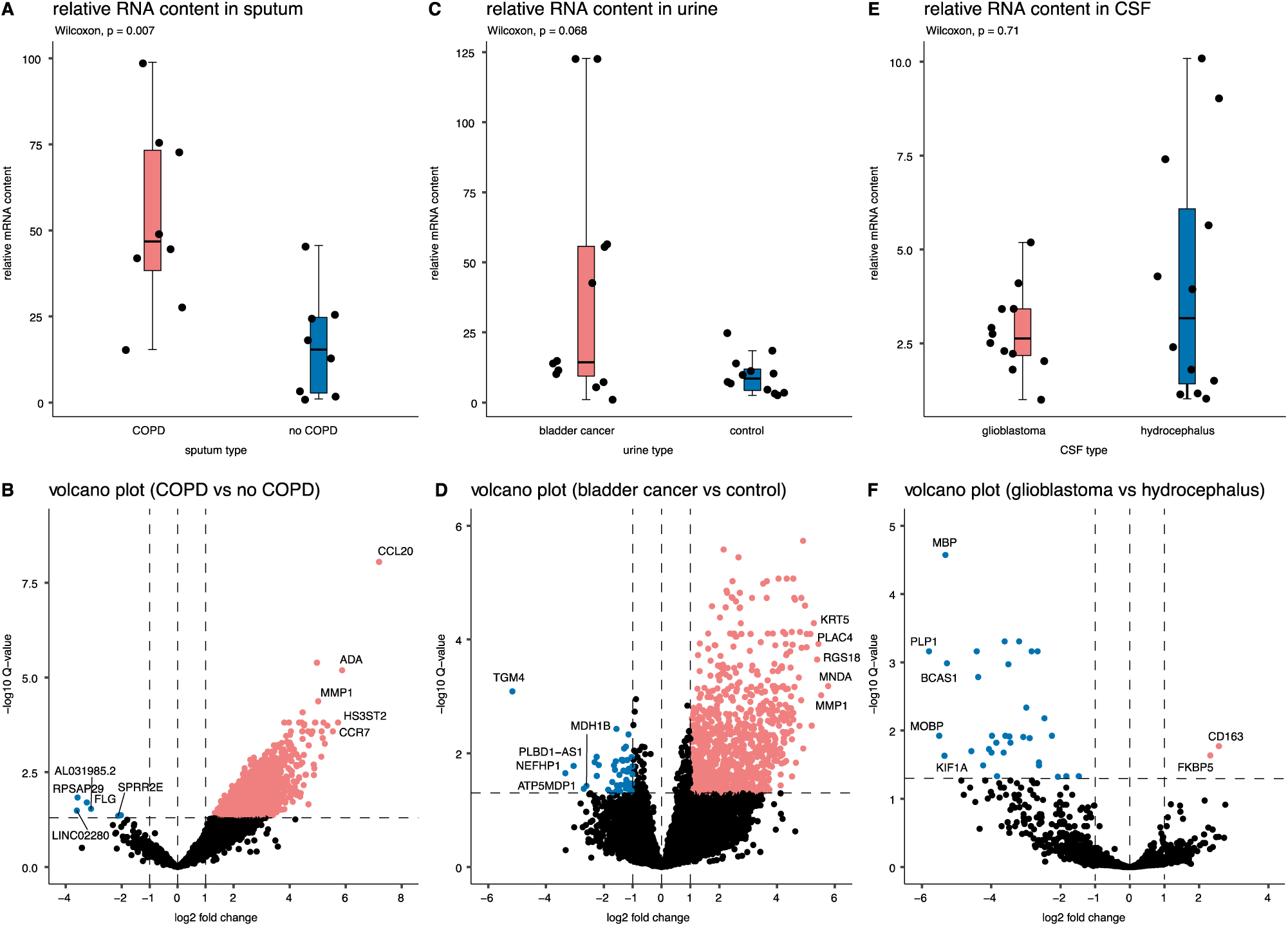
Relative RNA concentration and volcano plot in case/control cohorts. Top: Boxplots of relative mRNA content, bottom: Volcano plots of differentially expressed mRNAs (q<0.05; pink up; blue down in patient vs. control) with labeling of up to 5 most differential genes. (A) Sputum from COPD patients (n = 8) compared to sputum from healthy donors (n = 8; Wilcoxon rank test, two-sided, p = 0.007); (B) 5513 and 6 mRNAs up and down, respectively in COPD samples. (C) Urine from bladder cancer patients (n = 12) compared to urine from healthy donors (n = 12; Wilcoxon signed-rank test, two-sided, p = 0.068). (D) 529 and 9 mRNAs up and down, respectively in bladder cancer samples. (E) CSF from glioblastoma cancer patients (n = 12) compared to CSF from hydrocephalus patients (n = 12); Wilcoxon signed-rank test, two-sided, p = 0.71); (D) 2 and 33 mRNAs up and down, respectively in glioblastoma samples

In contrast to COPD, the relative RNA content is comparable in urine from bladder cancer patients and healthy volunteers, in CSF from glioblastoma patients and hydrocephalus patient, and in saliva from diabetes patients and healthy volunteers (Fig. 7C/E, Supplementary Fig. 10). A higher RNA yield in CSF from glioblastoma patients compared to CSF from healthy controls has been reported by Saugstad et al.^41^, however the collection method of CSF differed between both groups and it is therefore not possible to assess whether the reported difference in RNA yield between both groups is due to the different CSF collection site (lumbar puncture versus craniotomy) or due to the neurological disease. In urine from patients with a muscle invaded bladder cancer, 529 mRNAs and 9 mRNAs were significantly upregulated and downregulated, respectively, compared to urine from healthy volunteers (Fig. 7D). Some of the upregulated mRNAs, such as MDK, SLC2A1, GPRC5A, KRT17 and KRT5, have been reported in urine and were suggested as biomarker for the accurate detection and classification of bladder cancer^42–45^. In CSF from glioblastoma patients, only 2 mRNAs are significantly upregulated compared to CSF from hydrocephalus patients. CD163, one of the upregulated genes in glioblastoma, has been linked with glioblastoma pathogenesis^46^. In saliva from diabetes patients and saliva from healthy volunteers, no differentially expressed genes could be identified. A list with differentially expressed genes in all case/control cohorts can be found in Supplementary Data 5.

Differential abundance analysis was performed for circular RNAs as well, but in none of the case/control cohorts differentially abundant circRNAs could be detected (data not shown). As circular RNAs can only be identified based on their backsplice junction, the read coverage is generally (too) low for biomarker discovery based on mRNA capture sequencing data. When applying a similar strategy for mRNAs by looking at the reads of only one “linear only” junction per gene (outside every detected back-splice junction) a significantly lower number of differentially abundant mRNAs were detected (sputum: 13 out of 5519 mRNAs; urine: 0 out of 538 mRNAs; CSF: 0 out of 35 mRNAs). These results strongly suggest that a dedicated circRNA enrichment strategies may be needed to assess circRNA biomarker potential.

To validate the identification of the 10 most abundant circRNAs detected by mRNA capture sequencing in sputum, an orthogonal validation by RT-qPCR of the backsplice sequence region was performed. For 9 of the 10 circRNAs, the RNA-sequencing results could be validated. (Supplementary Fig. 9C)

## Discussion

By applying two complementary RNA-sequencing technologies on 20 different biofluids, we assembled the most comprehensive human biofluid transcriptome, covering small RNAs, mRNAs and circRNAs. Until now, most efforts to investigate and compare the RNA content within biofluids focused on small RNA sequencing, most likely because of technical limitations and unawareness of the abundance of extracellular mRNA (fragments)^5–7,9,12,13^.

The availability of both small RNA sequencing data and mRNA data allows a more in-depth characterization of the human transcriptome in biofluids. To our knowledge, this is the first study reporting on the mRNA content, generated through a dedicated mRNA enrichment sequencing method, in tear fluid, amniotic fluid, aqueous humor, bile, bronchial lavage fluid, gastric fluid, saliva, seminal plasma, synovial fluid, sweat and urine. Selected mRNAs were previously studied by means of RT-qPCR in amniotic fluid^14^, pancreatic cyst fluid^15,18^, seminal plasma^16^, sputum^17^, stool^19^ and in extracellular vesicles isolated from cell-free urine^20^. In saliva, selected mRNAs were detected using microarrays^47^. We have demonstrated that it is technically feasible to generate mRNA data from low input biofluid samples. This is expected to accelerate biomarker research in these fluids. Further efforts to profile and share the mRNA and circRNA content in larger sample cohorts of biofluids, comparable to the exRNA Atlas Resource for small RNAs, are necessary to move this scientific field forward.^8^

Our small RNA results confirm previous studies observing high miRNA concentration in tears^13^, low mapping rates in CSF^5,48^ and low miRNA concentration in cell-free urine^12^. A direct comparison of the absolute number of detected miRNAs, mRNAs and circRNAs detected per sample in our study with the numbers in published literature is hampered by the fact that the absolute read count is dependent on the input volume of the biofluids, the RNA isolation kit and library preparation method used, the sequencing depth and data-analysis settings (e.g. mapping without mismatches, filtering of the data). In addition, different pre-analytical variables when preparing the biofluid samples may also affect the sequencing results. However, on a higher level, we can look into the most abundant miRNAs detected in specific biofluids. The majority of the 10 most abundant miRNAs detected in 9 specific biofluids reported by Godoy et al. are also detected amongst the most abundant miRNAs in the samples from the discovery cohort (Supplementary Data 10)^5^.

We compared the mRNA results of the discovery cohort with these of the case/control cohorts. Mapping rates for samples in the discovery cohort are in the same range for saliva, sputum and seminal plasma. The mapping rates for CSF and urine are about 15% higher in the case/control cohorts compared to the discovery cohort. These differences may be due to different pre-analytical variables between both cohorts (collection tube, centrifugation speed and the portion of urine collected (Supplementary Fig. 3A; Supplemental material and methods).

In the discovery cohort on average 53% of all small RNA reads in saliva can be traced to bacteria, perfectly in line with the average of 45.5% reads mapping to bacteria reported by Yeri et al.^6^ Aqueous humor and CSF, although collected in a sterile setting and presumed to be sterile, contain up to 11% of reads mapping to bacteria, in line with previous studies^5,48^. However, bacterial cultures of our two CSF samples were negative. As both CSF and aqueous humor display a very low relative RNA content, the exogenous sequences may represent bacterial contaminants introduced during the sample processing workflow. Contaminants can derive from contaminated spin columns used during RNA purification^32^, enzymes produced in microorganisms ^49^, or various environmental sources^50^. Such contaminant signals are likely underrepresented in samples with high concentration of endogenous exRNAs.

Although we collected a broad range of biofluids, only two samples per biofluid were studied, limiting our ability to assess donor variability. The input volume for the RNA isolations in all biofluids was set to 200 µL and a volume-based comparison of the RNA content was made among the biofluids. We did not explore if higher input volumes would result in higher RNA yields in biofluids where this could have been possible (e.g. urine). We also note that the results in Table 1 are impacted by biofluid input volume in the RNA purification, RNA input in the sequencing library prep, and the sequencing depth.

Biofluid data normalization with synthetic spike-in controls is a unique and powerful approach and reflects more accurately the biological situation compared to classic normalization approaches where global differences on overall abundance are neutralized. For instance, the relative mRNA concentration in sputum from COPD patients is higher than in sputum from healthy donors. Typically, RNA sequencing data is subsampled or normalized based on the library size before performing a differential expression analysis, resulting in an artificially more balanced volcano plot, an overcorrection of the biological situation and a loss of information, which is not the case when the data is normalized based on spike-in controls.

Our results highlighting tissues and cell types that contribute more specifically to a certain biofluid compared to the other biofluids (Fig. 6A) can be used as a roadmap to formulate hypotheses when initiating biomarker research. Not surprisingly, the RNA signal from prostate is reflected in urine and seminal plasma. Both fluids can be collected in a non-invasive way and may be of value to investigate further in prostate cancer patients. Of interest, the mRNA concentration in seminal plasma is 1000-fold higher than in urine and seminal plasma contains more unique mRNAs compared to urine, suggesting that the biomarker potential of seminal plasma is higher. However, one should also be cautious in interpreting the tissue enrichment results: while the RNA signal of breast seems relatively enriched in sweat, this biofluid has the lowest RNA concentration. The limited number of detected mRNAs in sweat show overlap with mRNAs related to secretion (MCL1 gene, SCGB2A2 gene, SCGB1D2 gene) that also appear as markers in breast tissue.

The pancreatic tissue RNA signal appears to be enriched in pancreatic cyst fluid and a different cell type composition is observed when both samples are deconvoluted using single cell RNA sequencing data of pancreatic cell types (Fig. 6B). Pancreatic cyst fluid was collected in these donors to investigate a cystic lesion in the pancreas. The routine cytological analysis of these fluid samples was inconclusive at the moment of sample collection. By following up both patients, we discovered that the first patient developed a walled off necrosis collection after necrotizing pancreatitis. The incipient high fraction of activated stellate cells in the first cyst fluid sample may have been an indication pointing towards the inflammation and necrosis that finally occurred. The second patient was diagnosed with a side-branch intra papillary mucinous neoplasia, probably reflected by the relative high fraction of acinar cells. Pancreatic cysts are often detected on abdominal imaging, resulting in a diagnostic and treatment dilemma. Furthermore, pancreatic cysts represent a broad group of lesions, ranging from benign to malignant entities. The main challenge in their management is to accurately predict the malignant potential and to determine the risk to benefit of a surgical resection^51^. Our results show that the cellular contribution to the RNA content of pancreatic cyst fluids can be estimated through deconvolution and that these results may be associated with clinical phenotypes. Larger cohorts are necessary to investigate the clinical potential of this approach and pancreatic tumor cells may also need to be added to the reference set with single cell RNA sequencing data to improve the accuracy of the prediction.

In addition to linear mRNA transcripts, we also explored the circular RNA content in biofluids. CircRNAs are a growing class of non-coding RNAs and a promising RNA biotype to investigate in the liquid biopsy setting, as they are presumed to be less prone to degradation compared to linear forms^52^. The circRNA fraction in tissues has previously been reported and is in line with our findings^53^. In our study, we demonstrated that for genes that produce both circRNAs and linear mRNAs, the circRNAs are more abundant than the linear forms in biofluids. Further assessment of the biomarker potential of circRNAs in biofluids require dedicated library preparation methods with circRNA enrichment.

In conclusion, The Human Biofluid RNA Atlas provides a systematic and comprehensive comparison of the extracellular RNA content in 20 different human biofluids. The results presented here may serve as a valuable resource for future biomarker studies.

## Material and methods

### Donor material, collection and biofluid preparation procedure

Sample collection for the discovery cohort and sputum collection for the case/control cohort was approved by the ethics committee of Ghent University Hospital, Ghent, Belgium (no. B670201734450) and written informed consent was obtained from all donors according to the Helsinki declaration. Breast milk, colostrum, plasma, serum, sputum, seminal plasma, sweat, stool, tears and urine were obtained in healthy volunteers. All other biofluids were collected from non-oncological patients.

The collection of two case series of each 12 cases and 12 control samples was approved by the Masaryk Memorial Cancer Institute, Brno, Czech Republic (no. 14-08-27-01 and no. MOU190814). Urine was collected in healthy donors and muscle-invasive bladder cancer patients; CSF was collected in hydrocephalus patients and glioblastoma patients.

Collection of saliva samples in 12 healthy donors and in patients with diabetes mellitus for the case/control cohort was approved by the ethics committee of the Medical University of Vienna, Vienna, Austria (no. 2197/2015). Written informed consent was obtained from all donors. The demographic and clinical patient information is provided in Supplementary Table 1. Detailed information on the sample collection per biofluid is provided in Supplementary Note 2. All samples, except tear fluid, plasma and serum, were centrifuged at 2000 g (rcf) for 10 minutes without brake at room temperature. All samples were processed within 2 hours after collection. The cell-free supernatant was carefully pipetted into 2 mL LoBind tubes (Eppendorf LoBind microcentrifuge tubes, Z666556-250EA) and stored at −80 °C.

### RNA isolation and gDNA removal

#### RNA isolation from all biofluids, except tears

In the discovery cohort, two RNA isolations per biofluid and per sample were simultaneously performed by two researchers (E.V.E. and E.H.). In the end, RNA obtained from both RNA isolations was pooled per biofluid and per sample and this pooled RNA was used as starting material for both library preparations. Hence, small RNA and mRNA capture sequencing on the discovery cohort were performed on the same batch of RNA. In the case/control cohorts, one RNA isolation was performed per sample and the RNA was used as starting material for mRNA capture sequencing.

RNA was isolated with the miRNeasy Serum/Plasma Kit (Qiagen, Hilden, Germany, 217184) according to the manufacturer’s instructions. An input volume of 200 µL was used for all samples, except for tear fluid, and total RNA was eluted in 12 µL of RNAse-free water. Tear fluid was collected with Schirmer strips and RNA was isolated directly from the strips (see further). Per 200 µL biofluid input volume, 2 µL Sequin spike-in controls (Garvan Institute of Medical Research) and 2 µl RNA extraction Control (RC) spike-ins (Integrated DNA Technologies)^54^ were added to the lysate for TruSeq RNA Exome Library Prep sequencing and TruSeq Small RNA Library Prep sequencing, respectively. Details on the spike-in controls are available in the Supplementary Note 1.

Briefly, 2 µl External RNA Control Consortium (ERCC) spike-in controls (ThermoFisher Scientific, Waltham, MA, USA, 4456740), 2 µl Library Prep Control (LP) spike-ins (Integrated DNA Technologies)^55^, 1 µl HL-dsDNase and 1.6 µl reaction buffer were added to 12 µl RNA eluate, and incubated for 10 min at 37 °C, followed by 5 min at 55 °C. Per biofluid and per donor the RNA after gDNA removal was pooled. RNA was stored at −80 °C and only thawed on ice immediately before the start of the library prep. Multiple freeze/thaw cycles did not occur.

#### RNA isolation from tear fluid

Tear fluid was collected in 8 healthy donors with Schirmer strips (2 strips per eye per donor), as previously described^56,57^. RNA was isolated within two hours after tear collection with the miRNeasy Serum/Plasma Kit (Qiagen, Hilden, Germany, 217184), starting from one 2 mL tube containing each 4 Schirmer strips. The same reagent volumes as suggested by the manufacturer for a 200 µL input volume were used. Throughout the RNA isolation protocol, the two final RNA samples each result from 4 tear fluid samples (each containing the 4 strips of a single donor) that were pooled in a two-step method. First, the upper aqueous phase of two tear fluid samples was put together (in step 8 of the RNA isolation protocol). Second, the RNA eluate of these two samples was pooled into the final RNA that was used as input for the library prep (in step 15 of the RNA isolation protocol).

### TruSeq RNA Exome library prep sequencing

Messenger RNA capture based libraries were prepared starting from 8.5 µL DNase treated and spike-in supplemented RNA eluate using the TruSeq RNA Exome Library Prep Kit (Illumina, San Diego, CA, USA). Each sample underwent individual enrichment according to the manufacturer’s protocol. The quality and yield of the prepared libraries were assessed using a high sensitivity Small DNA Fragment Analysis Kit (Agilent Technologies, Santa Clara, CA, USA) according to manufacturer’s instructions. The libraries were quantified using qPCR with the KAPA Library Quantification Kit (Roche Diagnostics, Diegem, Belgium, KK4854) according to manufacturer’s instructions. Based on the qPCR results, equimolar library pools were prepared.

Paired-end sequencing was performed on a NextSeq 500 instrument using a high output v2 kit (Illumina, San Diego, CA, USA) with a read length of 75 nucleotides to an average sequencing depth of 11 million read pairs in the discovery cohort, 16.8 million read pairs in the sputum case/control cohorts, 15.4 million read pairs in the urine case/control cohort, 15 million read pairs in the CSF case/control cohort and 18.8 million read pairs in the saliva case/control cohort. Samples from the discovery cohort were randomly assigned over two pools and sequenced with a loading concentration of 1.2 pM (5% PhiX) and 1.6 pM (5% PhiX), respectively. Urine, CSF and saliva samples from the case/control cohorts were loaded in 3 separate runs at 2 pM (2% PhiX) and sputum samples from the case/control cohorts were loaded at 1.6 pM (5% PhiX).

### TruSeq Small RNA library prep sequencing

Small RNA libraries were prepared starting from 5 µL DNase treated and spike-in supplemented RNA eluate using a TruSeq Small RNA Library Prep Kit (Illumina, San Diego, CA, USA) according to the manufacturer’s protocol with two minor modifications(1). The RNA 3’ adapter (RA3) and the RNA 5’ adapter (RA5) were 4-fold diluted with RNase-free water(2) and the number of PCR cycles was increased to 16.

First, a volume-based pool of all 46 samples of the discovery cohort was sequenced. After PCR amplification, quality of libraries was assessed using a high sensitivity Small DNA Fragment Analysis Kit (Agilent Technologies, Santa Clara, CA, USA) according to manufacturer’s instructions. Size selection of the pooled samples was performed using 3% agarose dye-free marker H cassettes on a Pippin Prep (Sage Science, Beverly, MA, USA) following manufacturer’s instructions with a specified collection size range of 125–163 bp. Libraries were further purified and concentrated by ethanol precipitation, resuspended in 10 μl of 10 mM tris-HCl (pH = 8.5) and quantified using qPCR with the KAPA Library Quantification Kit (Roche Diagnostics, Diegem, Belgium, KK4854) according to manufacturer’s instructions. The pooled library was quality controlled via sequencing at a concentration of 1.7 pM with 35% PhiX on a NextSeq 500 using a mid-output v2 kit (single-end 75 nucleotides, Illumina, San Diego, CA, USA), resulting in an average sequencing depth of 1 million reads, ranging from 3341 reads to 14 million reads. Twenty-three samples with less than 200 000 reads were assigned to a low concentrated pool, 23 samples with more than 17 million reads were assigned to a highly concentrated pool. Based on the read numbers from the mid output run, two new equimolar pools were prepared, purified and quantified as described higher. Both re-pooled libraries were then sequenced at a final concentration of 1.7 pM with 25% PhiX on a NextSeq 500 using a high output v2 kit (single-end, 75 nucleotides, Illumina, San Diego, CA, USA), resulting in an average sequencing depth of 9 million reads (range 817 469 – 41.7 million reads).

### RT-qPCR

To validate findings observed in the RNA sequencing data, we performed a targeted mRNA and circRNA expression profiling with RT-qPCR for 8 differentially expressed mRNAs in sputum (COPD versus healthy control) and for the 10 most abundant circRNAs in sputum. As reference RNAs for normalization purposes, we selected Sequin spikes stably detected in all samples based on the available RNA sequencing data. The assays to measure mRNA, circRNA and Sequin spike expression were custom designed using primerXL^58^ (Supplementary Data 9) and purchased from Integrated DNA Technologies, Inc. (Coralville, USA).

For cDNA synthesis, 5 μl of total RNA was reverse transcribed using the iScript Advanced cDNA Synthesis Kit (BioRad, USA) in a 10 µL volume. 5 µL of cDNA was pre-amplified in a 12-cycle PCR reaction using the Sso Advanced PreAmp Supermix (Bio-Rad, USA) in a 50 µL reaction. Pre-amplified cDNA was diluted (1:8) and 2 µL was used as input for a 45-cycle qPCR reaction, quantifying 8 mRNAs and 10 circRNAs of interest with the SsoAdvanced™ Universal SYBR Green Supermix (BioRad, USA). All reactions were performed in 384-well plates on the LightCycler480 instrument (Roche) in a 5 µL reaction volume using 250 nM primer concentrations. Cq-values were determined with the LightCycler^®^480 Software (release 1.5.0, Roche) with the “Abs Quant/2nd Derivative Max” method.

The geNorm analysis to select the optimal number of reference targets was performed using Biogazelle’s qbase+ software (www.qbaseplus.com) using log2-transformed RNA count data. We observed medium reference target stability (average geNorm M ≤ 1.0) with an optimal number of reference targets in this experimental situation of two (geNorm V < 0.15 when comparing a normalization factor based on the two or three most stable targets). As such, the optimal normalization factor can be calculated as the geometric mean of reference targets R2_150 and R2_65. These Sequin spike RNAs were considered as reference RNAs.

### Data analysis

#### Processing TruSeq RNA Exome sequencing data

Read quality was assessed by running FastQC (v0.11.5) on the FASTQ files and reads shorter than 35 nucleotides and with a quality (phred) score < 30 were removed. The reads were mapped with STAR (v2.6.0). Mapped reads were annotated by matching genomic coordinates of each read with genomic locations of mRNAs (obtained from UCSC GRCh38/hg38 and Ensembl, v91) or by matching the spike-in sequences. Picard (v2.18.5) was used for duplicate removal. HTSeq (v0.9.1) was used for quantification of PCR deduplicated reads. A cut-off for filtering noisy genes was set based on historic data to remove noisy genes. Using a threshold of 4 counts, at least 95% of the single positive replicate values are filtered out. A table with the read count of mRNAs per sample is provided in Supplementary Data 6.

#### Processing TruSeq Small RNA sequencing data

Adaptor trimming was performed using Cutadapt (v1.8.1) with a maximum error rate of 0.15. Reads shorter than 15 nts and those in which no adaptor was found were discarded. For quality control the FASTX-Toolkit (v0.0.14) was used, a minimum quality score of 20 in at least 80% of nucleotides was applied as a cutoff. The reads were mapped with Bowtie (v1.1.2) without allowing mismatches. Mapped reads were annotated by matching genomic coordinates of each read with genomic locations of miRNAs (obtained from miRBase, v22) and other small RNAs (obtained from UCSC GRCh38/hg38 and Ensembl, v91) or by matching the spike-in sequences. Reads assigned as “not annotated” represent uniquely mapped reads that could not be classified in one of the small RNA biotype groups. As for the mRNA data, genes with fewer than 4 counts were filtered out. A table with the read count of miRNAs per sample is provided in Supplementary Data 7.

#### Exogenous RNA characterization

The exogenous RNA content in the mRNA data was assessed using the MetaMap pipeline^59^. Briefly, all reads were mapped to the human reference genome (hg38) using STAR (v2.5.2)^60^. Unmapped reads were subsequently subjected to metagenomic classification using CLARK-S (v1.2.3)^61^. Reads were summed across all bacterial species.

The exogenous RNA content in the small RNA data was assessed using the exceRpt small RNA-seq pipeline (v4.6.2) in the Genboree workbench with default settings^62^. Briefly, after adapter trimming, read quality was assessed by FASTQC (v0.11.2). A minimum quality score of 20 in at least 80% of nucleotides was applied as cutoff. The minimum read length after adapter trimming was set to 18 nucleotides. Reads were first mapped to the custom spike-in sequences using Bowtie2 (v2.2.6), followed by mapping the unmapped reads with STAR (v2.4.2a) to UniVec contaminants and human ribosomal (rRNA) sequences to exclude them before mapping (also with STAR) to the following databases: miRbase (v21), gtRNAdb, piRNABank, GeneCode version 24 (hg38) and circBase (version last updated in July 2017). A single mismatch was allowed during mapping to the human genome. Unmapped reads were then mapped with STAR to exogenous miRNAs and rRNAs. In the end, the remaining unmapped reads were mapped to the genomes of all sequenced species in Ensembl and NCBI. No mismatches were allowed during exogenous alignment. Raw read counts obtained from the Genboree workbench were further analyzed in R (v3.5.1) making use of tidyverse (v1.2.1).

#### Circular RNA detection and circular/linear ratio determination

Only TruSeq RNA Exome reads passing quality control (base calling accuracy of ≥ 99% in at least 80% of the nucleotides in both mates of a pair) were included in this analysis. Clumpify dedupe (v38.26) was used to remove duplicates in paired-end mode (2 allowed substitutions, kmer size of 31 and 20 passes). We used a two-step mapping strategy to identify forward splice (further referred to as linear) junction reads and backsplice junction reads. First, reads were aligned with TopHat2 (v2.1.0) to the GRCh38/hg38 reference genome (Ensembl, v91). Micro-exons were included, a minimum anchor length of 6 nucleotides was required, and up to two mismatches in the anchor region were allowed. The resulting output contains linear junction information. Secondly, unmapped reads from the first mapping strategy were realigned with TopHat2 (v2.1.0) to the same reference, but this time with the fusion search option that can align reads to potential fusion transcripts. Processing the fusion search output with CIRCexplorer2 parse (v2.3.3) results in backsplice junction information. Junction read counts obtained with the mapping strategies described above were used as a measure for the relative level of linear and circular RNA in each sample. Only genes with at least one detected backsplice junction were considered. Junctions that could be part of both linear and circular transcripts (ambiguous junctions) were filtered out. As there is currently no consensus on how to calculate the circular to linear ratio (CIRC/LIN), we decided to calculate the ratio in two different ways (Supplementary Fig. 8). The circRNA fraction is defined as 100*CIRC/(CIRC+LIN). The first method (referred to as “backsplice junction-level method”) zooms in on each particular backsplice junction. CIRC was defined as the backsplice junction read count of one particular backsplice junction. LIN was defined as the average read count of all junctions flanking the backsplice junction of interest. The second method (referred to as “gene-level method”) considers all backsplice junctions within a given gene. CIRC was defined as the average number of backsplice junction reads for a given gene. LIN was defined as the average number of linear junction reads for a given gene. For both methods, CIRC > 3 was used as a cut-off for filtering noisy backsplice junctions. To enable a comparison of the circular/linear genic ratios in biofluids with those of tissues, the mRNA capture sequencing FASTQ files of 16 cancerous tissue types (34 samples in total) were downloaded from the MiOncoCirc database (dbGaP Study Accession phs000673.v3.p1)^24^. A list with the downloaded samples is attached in Supplementary Table 2. A table with the read count of backsplice junctions per sample is provided in Supplementary Data 8.

#### Assessment of tissue and cell contribution to biofluid exRNA

Using total RNA-sequencing data from 27 normal human tissue types and 5 immune cell types from peripheral blood from the RNA Atlas^35^, we created gene sets containing marker genes for each individual entity (Supplementary Data 4). We removed redundant tissues and cell types from the original RNA Atlas (e.g. granulocytes and monocytes were present twice; brain was kept and specific brain sub-regions such as cerebellum, frontal cortex, occipital cortex and parietal cortex were removed) and we used genes where at least one tissue or cell type had expression values greater or equal to 1 TPM normalized counts. A gene was considered to be a marker if its abundance was at least 5 times higher in the most abundant sample compared to the others. For the final analysis, only tissues and cell types with at least 3 markers were included, resulting in 26 tissues and 5 immune cell types.

Gene abundance read counts from the biofluids were normalized using Sequin spikes as size factors in DESeq2 (v1.22.2). For all marker genes within each gene set, we computed the log2 fold changes between the median read count of a biofluid sample pair versus the median read count of all other biofluids. The median log2 fold change of all markers in a gene set was selected, followed by z-score transformation over all biofluids (Fig. 7). For visualization purposes, only tissues and cell types with a z-score ≥ |1| in at least one biofluid were used.

#### Cellular deconvolution of pancreatic cyst fluid samples

To build the reference matrix for the computational deconvolution of pancreatic cyst fluid samples, single cell RNA sequencing data of 10 pancreatic cell types^34^ was processed with the statistical programming language R (v3.6.0). For each gene, the mean count across all individual cells from each cell type was computed. Next, this reference matrix was normalized using the trimmed means of M values (TMM) with the edgeR package (v3.26.4)^63,64^. Limma-voom (v3.40.2)^65^ was used for subsequent differential gene expression analysis and those genes with an absolute fold change greater or equal to 2 and an adjusted p-value < 0.05 (Benjamini-Hochberg) were retained as markers^66^. Finally, using these markers and both the pancreatic cyst fluid samples and the reference matrix described above, the cell type proportions were obtained through computational deconvolution using non-negative least squares (nnls package; v1.4)^67,68^.

#### Differential expression analysis in case/control cohorts

Further processing of the count tables was done with R (v3.5.1) making use of tidyverse (v1.2.1). Gene expression read counts from the biofluids were normalized using Sequin spikes as size factors in DESeq2 (v1.20.0)^69^. To assess the biological signal in the case/control cohorts, we performed differential expression analysis between the patients and control groups using DESeq2 (v1.20.0). Genes were considered differentially expressed when the absolute log2 fold change > 1 and at q < 0.05.

### Data availability

The raw RNA-sequencing data have been deposited at the European Genome-phenome Archive (EGA) under accession number EGAS00001003917. All spike-normalized sequencing data can be readily explored in the interactive web-based application R2: Genomics analysis and visualization platform (http://r2.amc.nl), and via a dedicated accessible portal (http://r2platform.com/HumanBiofluidRNAAtlas). This portal allows the analysis and visualization of mRNA, circRNA and miRNA abundance, as illustrated in Supplementary Fig. 11. All samples can be used for correlation, principle component, and gene set enrichment analyses, and many more. All other data are available within the article and supplementary information.

### Code availability

The R scripts to reproduce the analyses and plots reported in this paper are available from the corresponding authors upon request.

## Supporting information

Description of Supplementary Files

## Acknowledgments

E.H. is a recipient of a grant of the Fund for Scientific Research Flanders (FWO). F.A.C. and A.M. are supported by Special Research Fund (BOF) scholarship of Ghent University (BOF.DOC.2017.0026.01, BOF.DOC.2019.0047.01). A.G. is senior clinical researcher of the Fund for Scientific Research Flanders (FWO; 1805718N). This research is partly funded by “RNA-MAGIC”, “LNCCA” Concerted Research Actions of Ghent University (BOF19/GOA/008, BOF16/GOA/023), “Kom Op Tegen Kanker” and by the Czech Ministry of Health grants (NV18-03-00398, NV18-03-00360). We thank Tim Mercer for providing the Sequin spikes.

The results published here are in part based upon data generated by the Clinical Sequencing Exploratory Research (CSER) consortium established by the NHGRI. Funding support was provided through cooperative agreements with the NHGRI and NCI through grant numbers U01 HG006508 (Exploring Cancer Medicine for Sarcoma and Rare Cancers). Information about CSER and the investigators and institutions who comprise the CSER consortium can be found at http://www.genome.gov/27546194.

## Contributions

J.V. and P.M. conceived and supervised the project; E.H., K.V., N.Y., E.V.E., J.N. designed and performed the experiments; E.H., A.M., J.A. and F.A.C. analyzed the data; L.S. and S.K. performed analysis using the MetaMap pipeline; G.S. and S.K. contributed technical support and resources; E.H., A.G., P.H., P.J., G.B., K.B., T.M., T.D., V.N., C.V.C., K.R., E.R., D.H., K.T., O.S., C.N. collected samples; S.L. designed RT-qPCR primers; E.H., P.M and J.V. wrote the paper; J.K. developed dedicated tools to analyze RNA atlas data and results and implemented them in the online portal R2. All authors contributed to manuscript editing and approved the final draft.

