## Supplementary Information for "Charting extracellular transcriptomes in The Human Biofluid RNA Atlas"

Hulstaert et al.

### SUPPLEMENTARY FIGURES

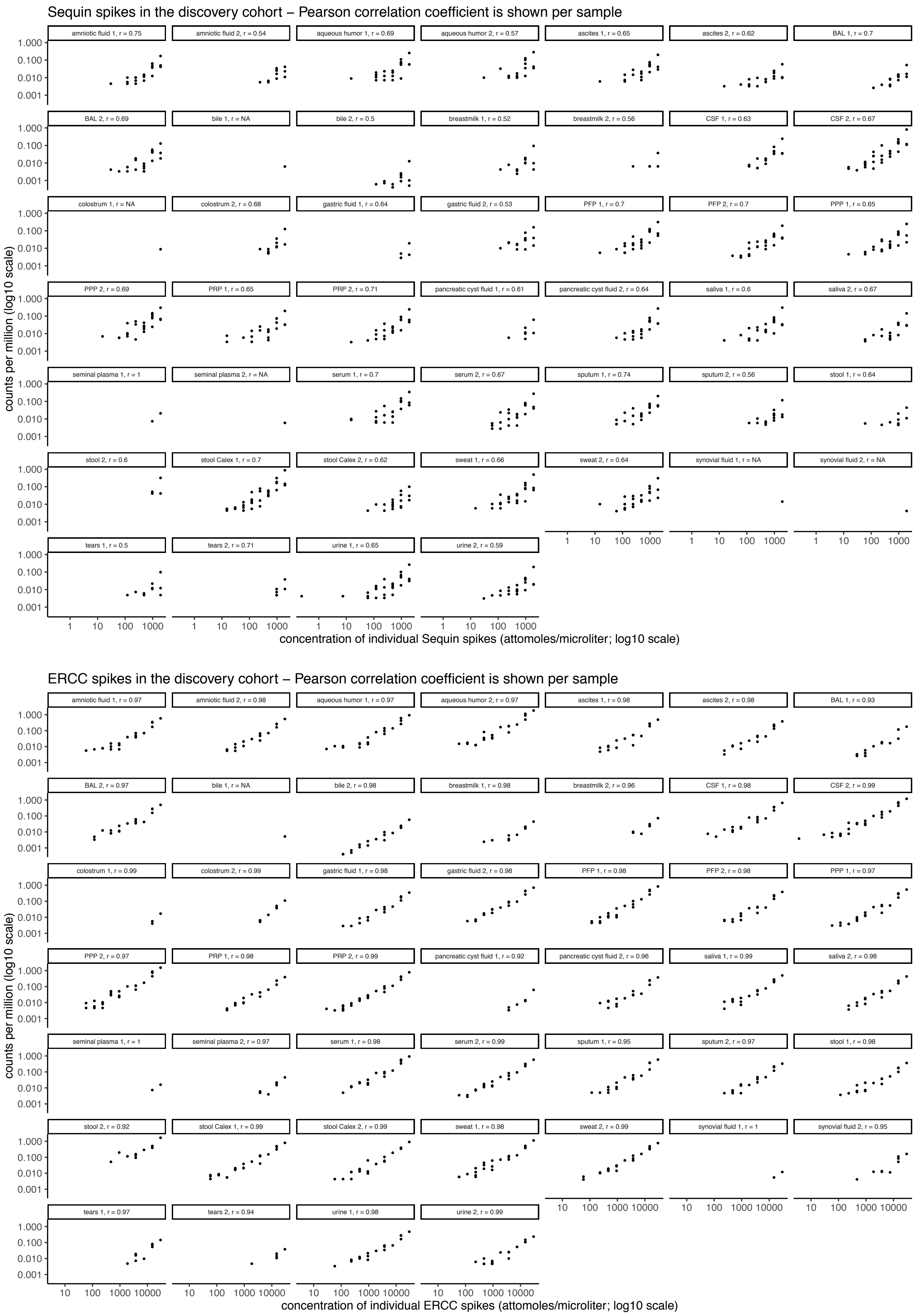

**Supplementary Figure 1. In the discovery cohort the expected and the observed relative quantities are correlated within each spike type**

Correlation plots per spike type and per sample are shown. The Pearson correlation coefficient is shown per sample.

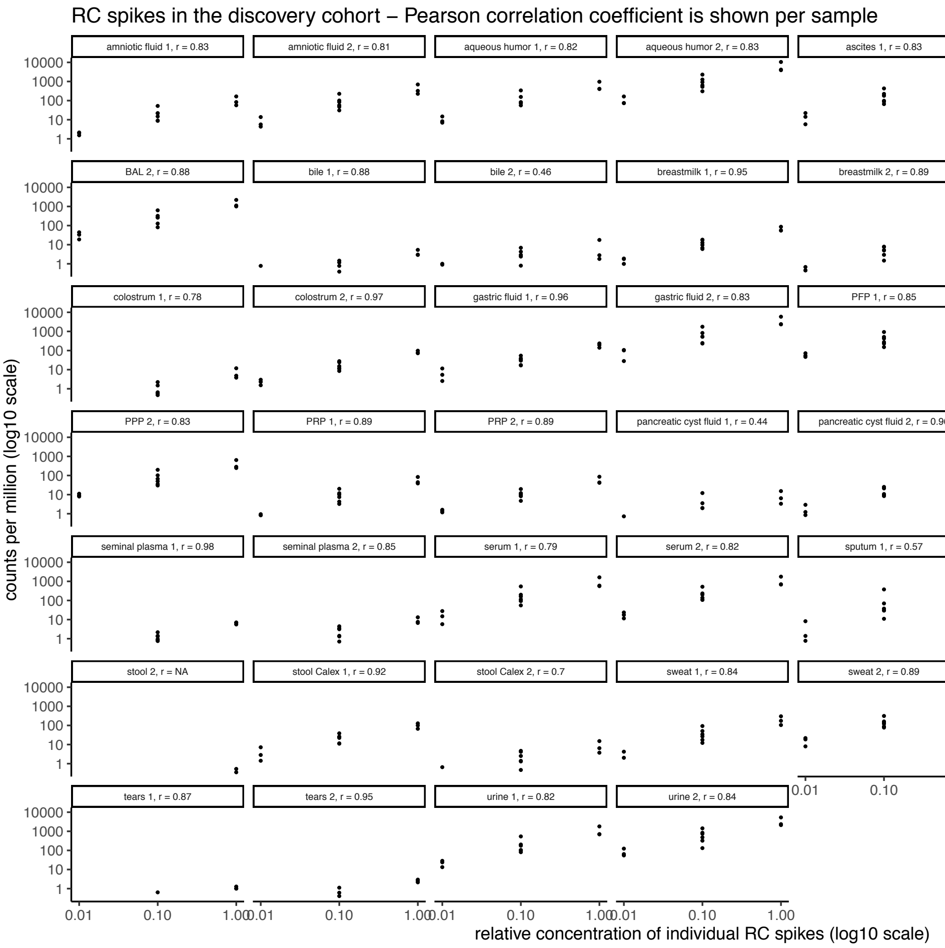

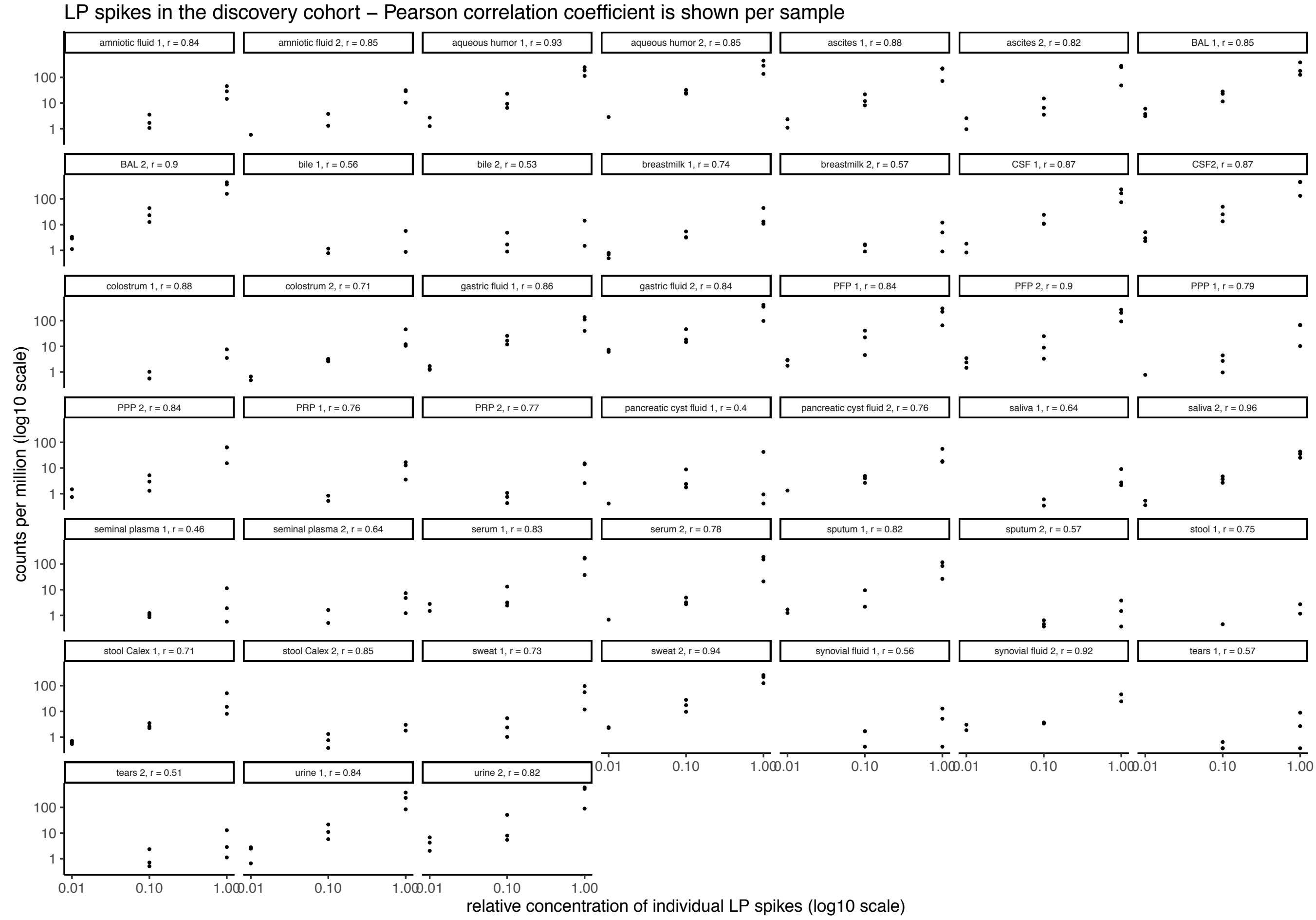

**Supplementary Figure 1. In the discovery cohort the expected and the observed relative quantities are correlated within each spike type**

Correlation plots per spike type and per sample are shown. The Pearson correlation coefficient is shown per sample.

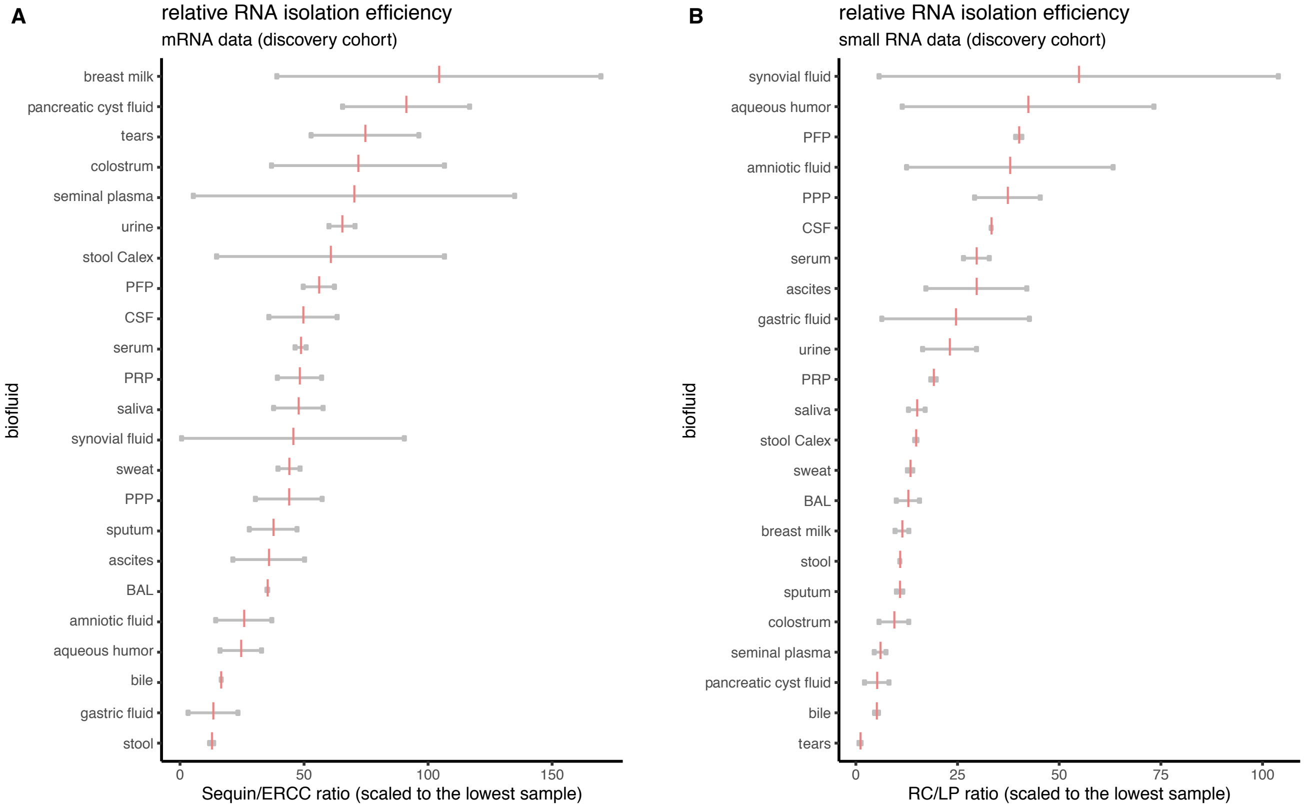

**Supplementary Figure 2. The relative RNA isolation efficiency in the discovery cohort varies across the different biofluids**

(A) The Sequin/ERCC ratio reflects the relative RNA isolation in the mRNA capture sequencing data and is plotted per biofluid. A minimal read count threshold is set on 15 reads per spike type. Each grey dot represents a sample. The pink vertical bar reflects the mean of both samples within a biofluid.

(B) The RC/LP ratio reflects the relative RNA isolation in the mRNA capture sequencing data and is plotted per biofluid. A minimal read count threshold is set on 15 reads per spike type. Each grey dot represents a sample. The pink vertical bar reflects the mean of both samples within a biofluid.

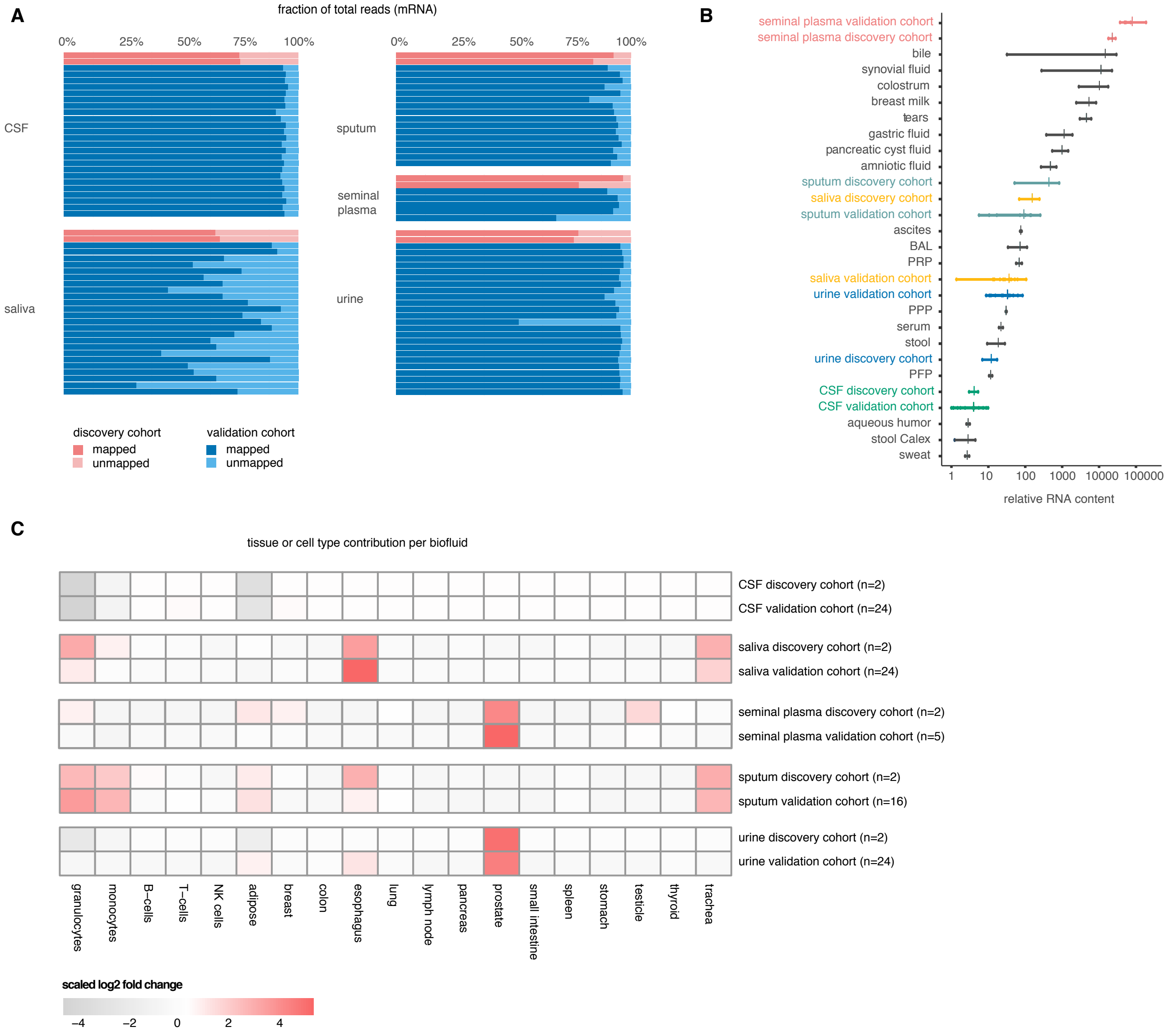

**Supplementary Figure 3. Mapping percentage, relative RNA content and tissue of origin analysis in additional CSF, saliva, sputum, seminal plasma and urine samples to illustrate the generalizability of the findings regarding the mRNA content observed in the discovery cohort**

(A) Percentage of the total read count mapping to the human transcriptome for CSF, saliva, sputum, seminal plasma and urine samples. Samples from the discovery cohort (n=2 per biofluid) are shown in pink. Samples from the validation cohort are shown in blue: CSF from 12 patients with hydrocephalus and 12 patients with glioblastoma, saliva from 12 healthy donors and 12 diabetes patients, sputum from 8 healthy donors and 8 COPD patients, seminal plasma from 5 healthy donors, urine from 12 healthy donors and 12 bladder cancer patients.

(B) Relative RNA concentration per biofluid; every dot represents the relative RNA concentration in one sample, every vertical mark indicates the mean per biofluid. Biofluids in black represent samples from the discovery cohort. For biofluids in color additional samples were analyzed: seminal plasma from 5 healthy donors, sputum from 8 healthy donors, saliva from 12 healthy donors, urine from 12 healthy donors and CSF from 12 patients with hydrocephalus.

(C) Assessment of the tissues of origin in CSF, saliva, seminal plasma, sputum and urine. Findings from the discovery cohort are validated in independent, additional samples. Heatmap showing tissues and cell types that contribute more specifically to a certain biofluid compared to the other biofluids. Rows depict the biofluids of the discovery cohort and the columns are the tissues or cell types for which markers were selected based on the RNA Atlas32. For visualization purposes, only tissues and cell types with a z-score transformed log2 fold change ≥ |1| in at least one biofluid are shown.

**Supplementary Figure 4. Correlation between samples of the same biofluid**

(A) Correlation plots of the mRNA data between two samples of the same biofluid are shown. The Pearson correlation coefficient is shown per biofluid.
(B) Correlation plots of the miRNA data between two samples of the same biofluid are shown. The Pearson correlation coefficient is plotted per biofluid.
BAL, bronchoalveolar lavage fluid; CSF, cerebrospinal fluid; HNSCC: head and neck squamous-cell carcinoma; PFP, platelet-free plasma; PPP, platelet-poor plasma; PRP, platelet-rich plasma

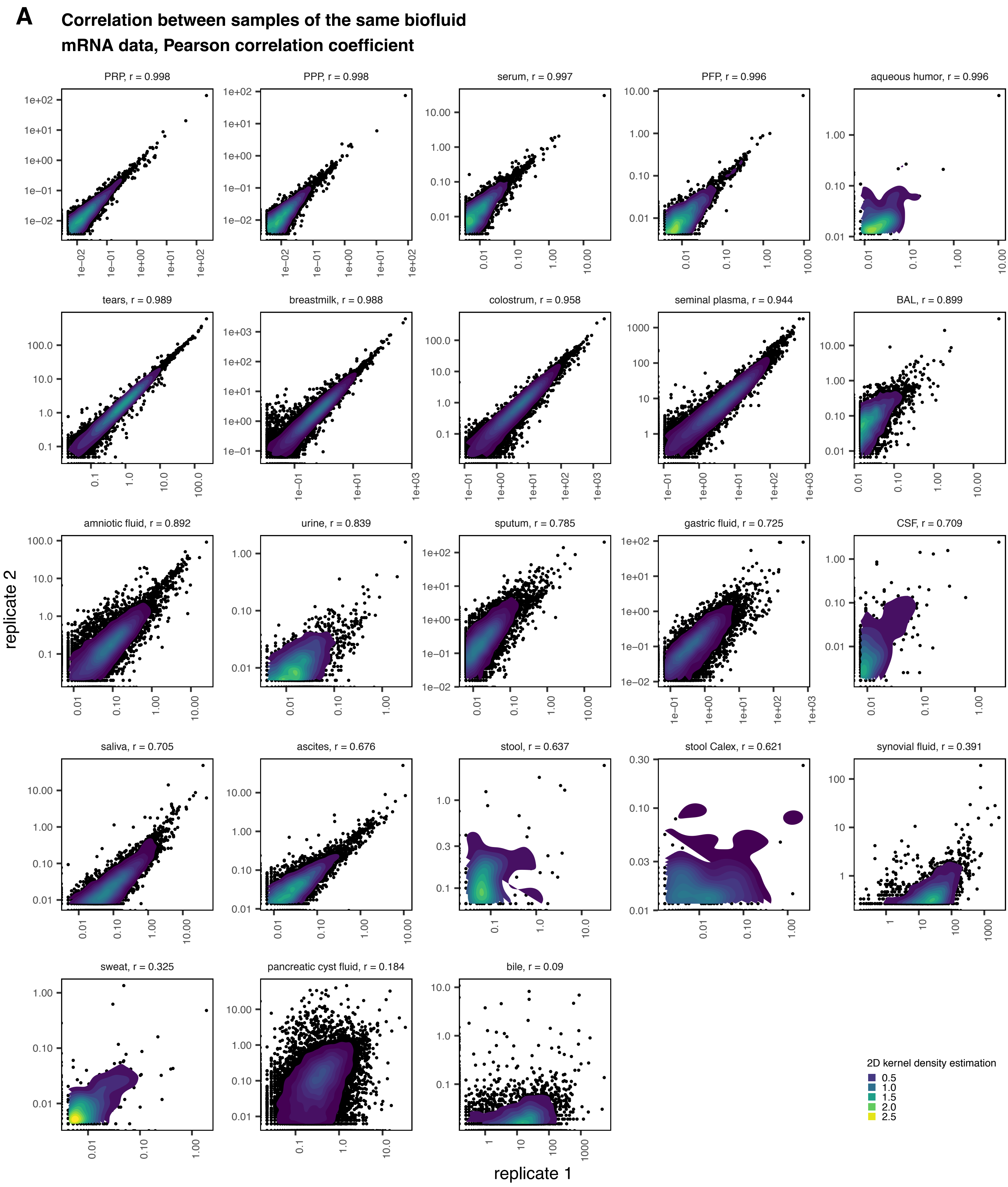

**Supplementary Figure 4. Correlation between samples of the same biofluid**

(A) Correlation plots of the mRNA data between two samples of the same biofluid are shown. The Pearson correlation coefficient is shown per biofluid.
(B) Correlation plots of the miRNA data between two samples of the same biofluid are shown. The Pearson correlation coefficient is plotted per biofluid.
BAL, bronchoalveolar lavage fluid; CSF, cerebrospinal fluid; HNSCC: head and neck squamous-cell carcinoma; PFP, platelet-free plasma; PPP, platelet-poor plasma; PRP, platelet-rich plasma

**B** **Correlation between samples of the same biofluid**
**miRNA data, Pearson correlation coefficient**

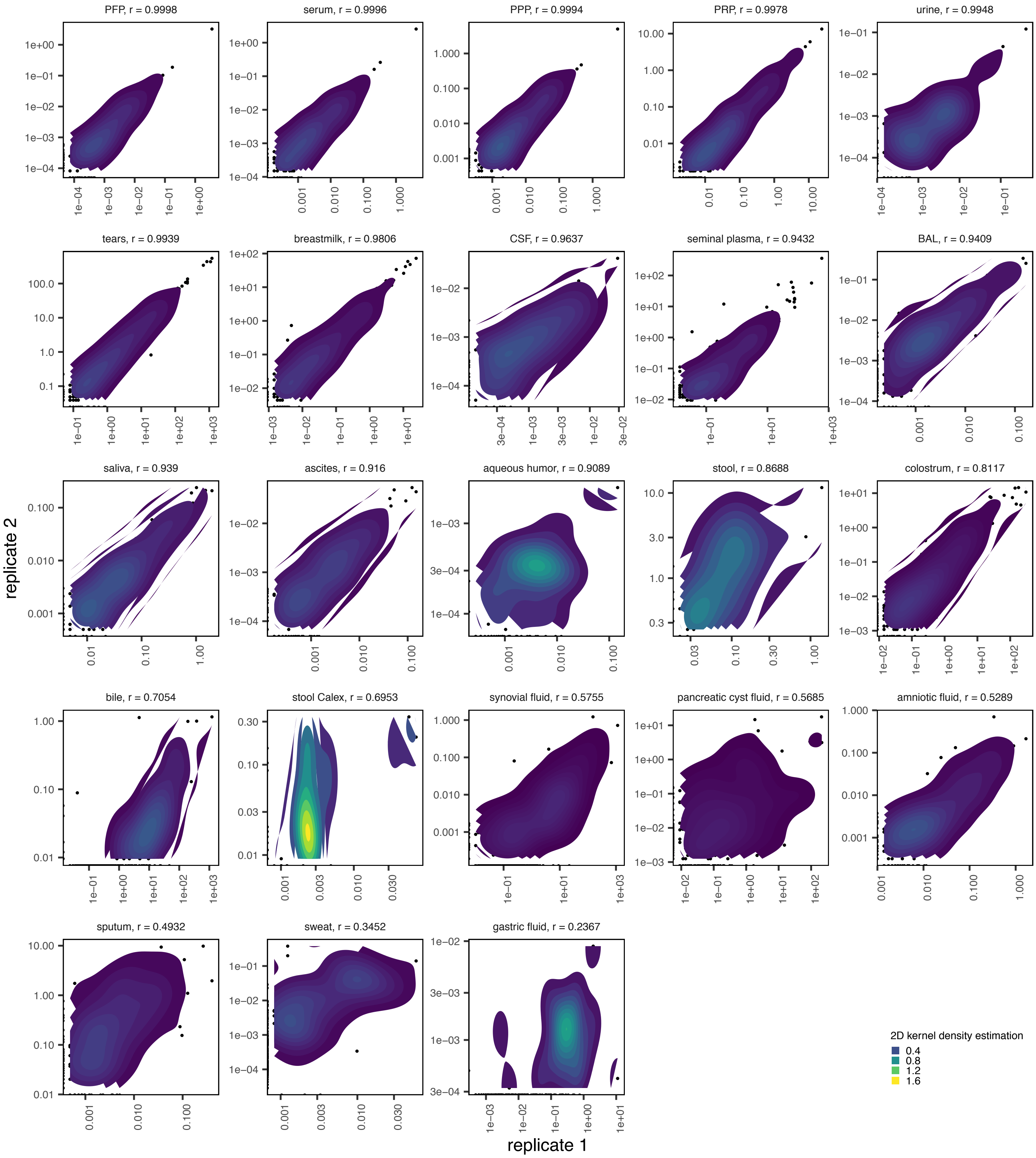

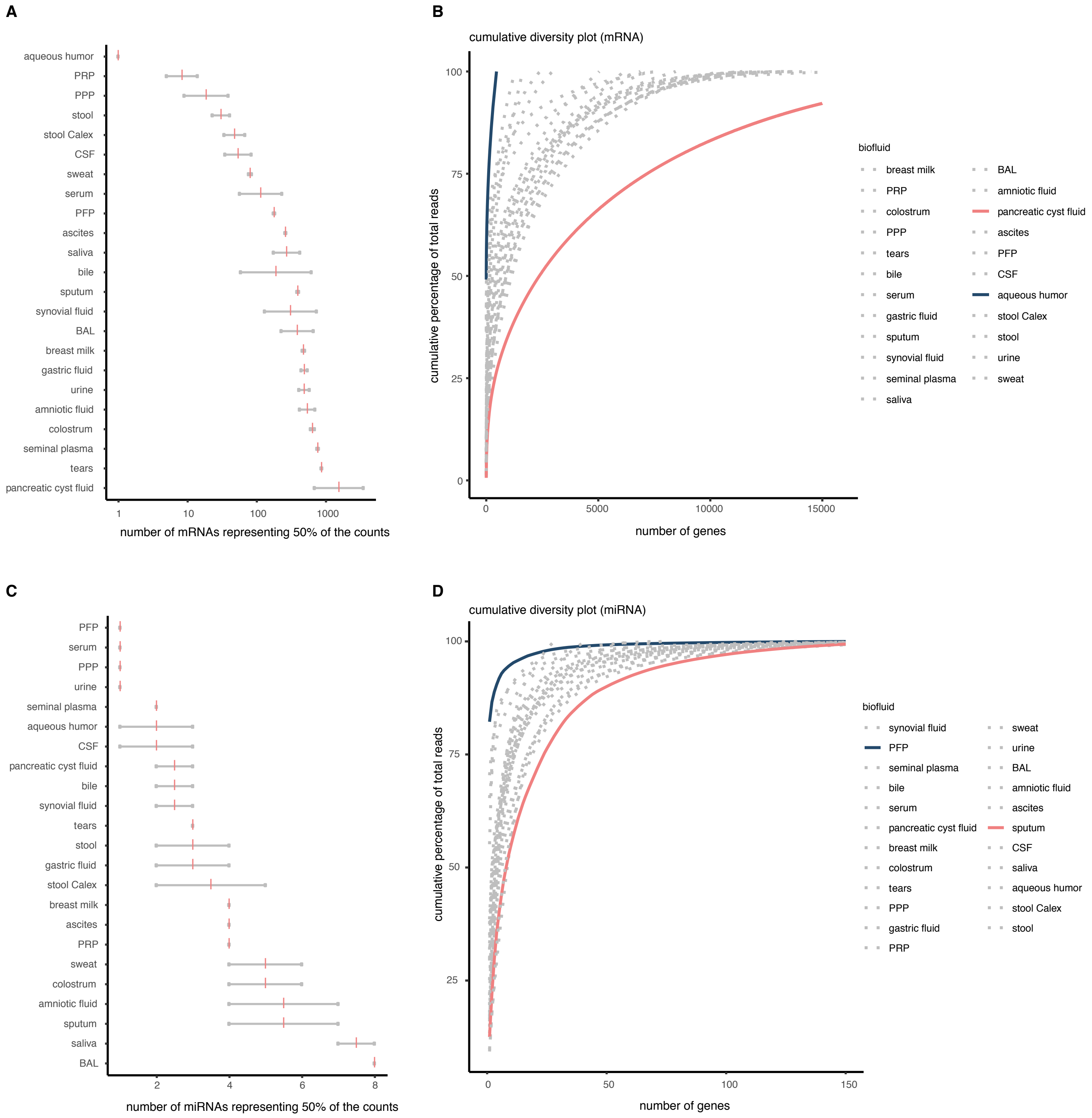

**Supplementary Figure 5. Additional RNA diversity metrics**

(A) The number of mRNAs representing 50% of the counts is shown per biofluid. Each gray dot represents a sample, the pink vertical bar represents the mean value within a biofluid.

(B) Cumulative diversity plot for the mRNA capture sequencing data.

(C) The number of miRNAs representing 50% of the counts is shown per biofluid. Each gray dot represents a sample, the pink vertical bar represents the mean value within a biofluid.

(D) Cumulative diversity plot for the miRNA data.

**A**

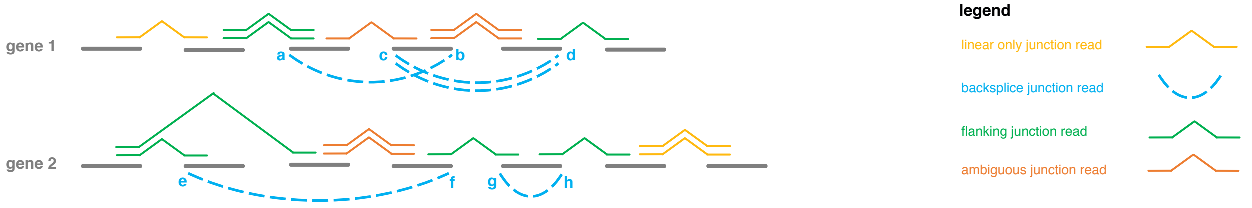

**B**

| | backsplice junction-level method | | gene-level method |
|---|---|---|---|
| gene 1 circ_a_b | CIRC = backsplice junction read = 1<br>LIN = average read count of all junctions flanking the backsplice junction of interest = $\frac{2}{1}$ + 0 = 2<br><br>**circRNA fraction = 33%** | gene 1 | CIRC = average number of backsplice junction reads for a given gene = $\frac{3}{2}$ = 1.5<br>LIN = average number of linear junction reads for a given gene = $\frac{1}{1}$ = 1<br><br>**circRNA fraction = 60%** |
| gene 2 circ_e_f | CIRC = backsplice junction read = 1<br>LIN = average read count of all junctions flanking the backsplice junction of interest = $\frac{\frac{2}{2} + \frac{1}{1}}{2}$ = 1<br>**circRNA fraction = 50%** | gene 2 | CIRC = average number of backsplice junction reads for a given gene = $\frac{2}{2}$ = 1<br>LIN = average number of linear junction reads for a given gene = $\frac{2}{1}$ = 2<br>**circRNA fraction = 33%** |

**Supplementary Figure 6. The circular RNA fraction was assessed using two different methods: the backsplice junction-level method and the gene-level method**

(A) Two genes and their different detected junction reads are shown to illustrate the terminology used throughout the manuscript. Junction reads that can only originate from linear transcripts (linear junction reads) are shown in yellow. Backsplice junction reads (shown in blue) can only be part of circular transcripts. Junction reads flanking a backsplice junction read are categorized as flanking junction read (shown in green). Junction reads that can be part of both linear and circular transcripts are referred to as ambiguous junction reads (shown in orange). Ambiguous junction reads are not taken into account for the further analyses.

(B) The circRNA fraction is defined as 100*CIRC/(CIRC+LIN). As an example the calculation of CIRC and LIN according to the backsplice junction-level method and the gene-level method is given below the figure.

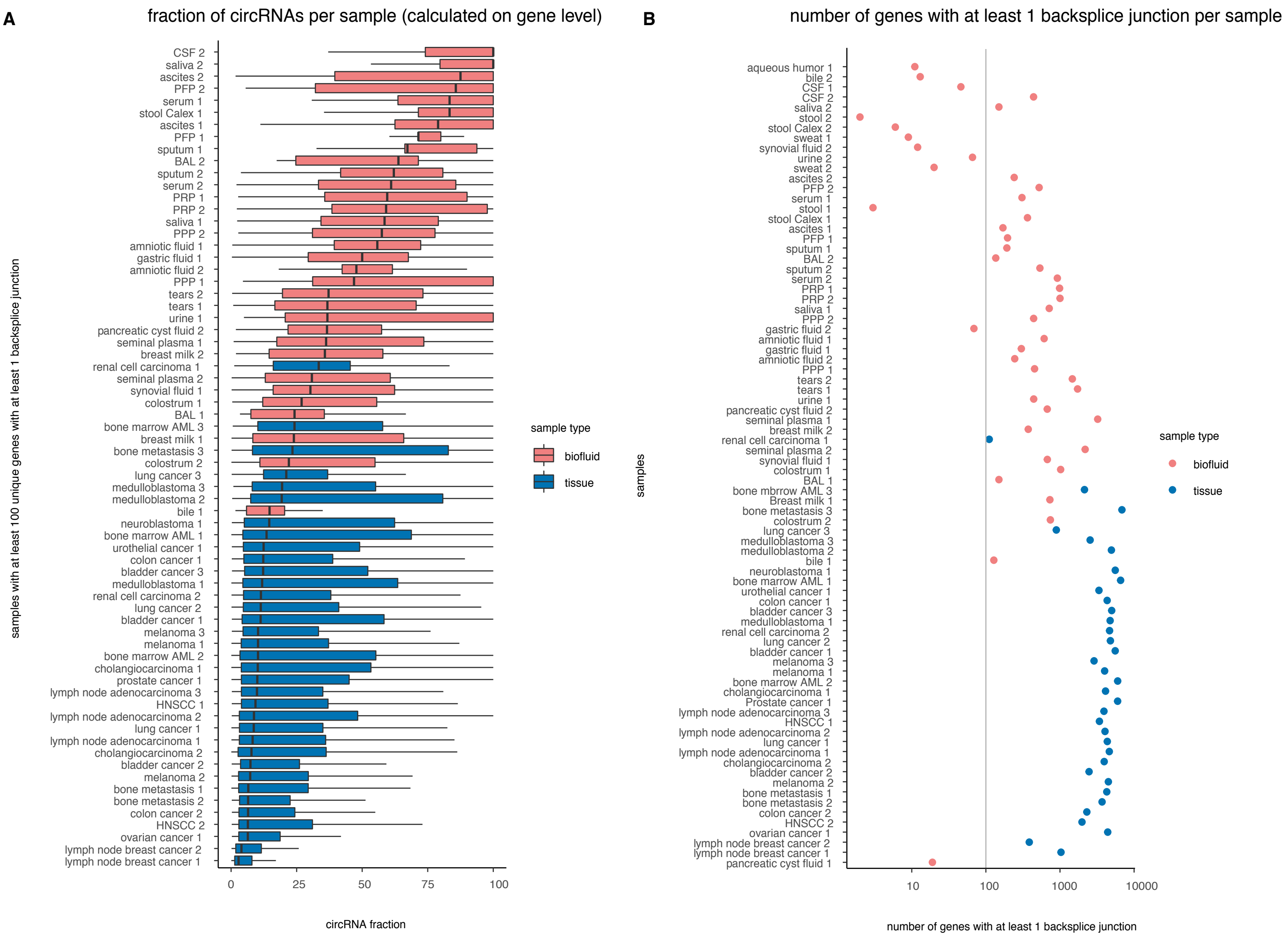

**Supplementary Figure 7. CircRNAs are enriched in biofluids compared to tissues**

(A) The circRNA fraction, calculated on gene level, is plotted per sample and is higher in cell-free biofluid samples than in tissue samples. Only samples with at least 100 backsplice junctions are plotted.

(B) The number of genes with at least 1 backsplice junction per sample is higher in tissue samples compared to biofluid samples, in line with the higher input concentration of RNA.

AML, acute myeloid leukemia; BAL, bronchoalveolar lavage fluid; CSF, cerebrospinal fluid; HNSCC: head and neck squamous-cell carcinoma; PFP, platelet-free plasma; PPP, platelet-poor plasma; PRP, platelet-rich plasma

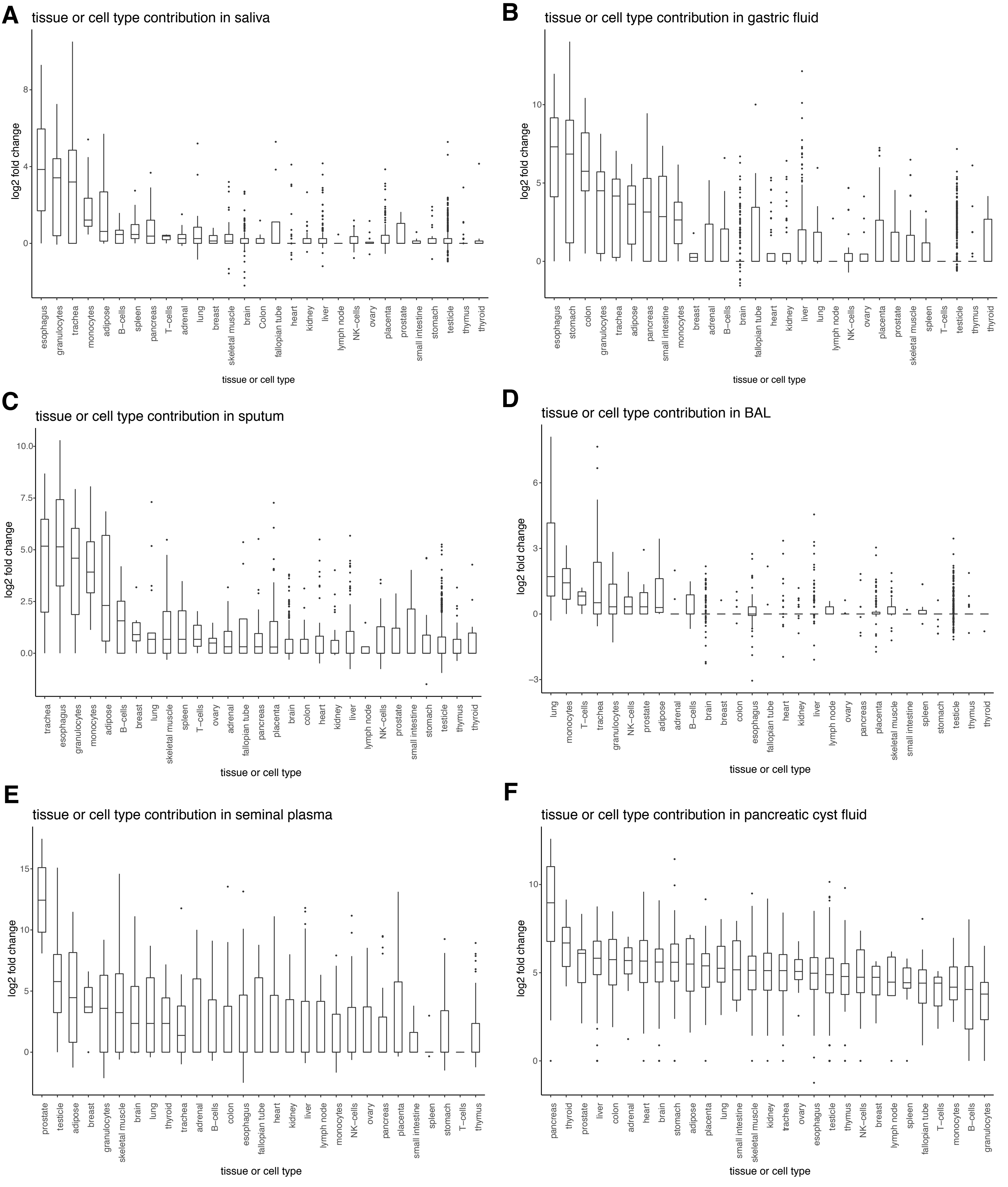

**Supplementary Figure 8. Tissue or cell type contribution in selected biofluids from the discovery cohort**

Boxplots showing the log2 fold change for a gene set with markers specific for a certain tissue or cell type. The log2 fold change is calculated between the median read count for both samples of 1 biofluid (indicated in the title of each plot) and the median read count of all other biofluids. The tissues or cell types for which markers were selected based on the RNA Atlas Project are shown on the x-axis.

(A) Tissue or cell type contribution in saliva
(B) Tissue or cell type contribution in gastric fluid
(C) Tissue or cell type contribution in sputum
(D) Tissue or cell type contribution in bronchoalveolar lavage fluid (BAL)
(E) Tissue or cell type contribution in seminal plasma
(F) Tissue or cell type contribution in pancreatic cyst fluid

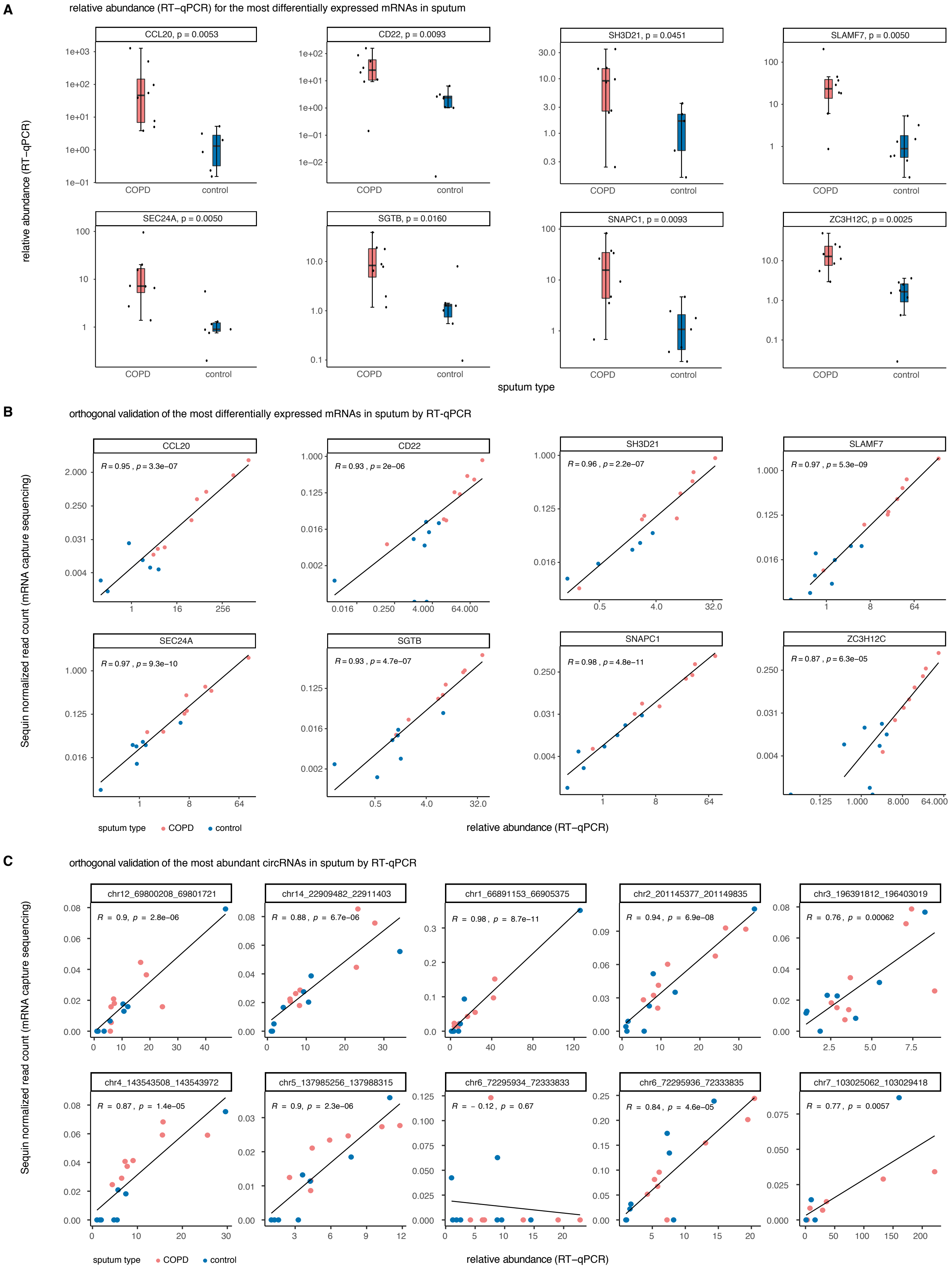
A
relative abundance (RT−qPCR) for the most differentially expressed mRNAs in sputum
CCL20, p = 0.0053
CD22, p = 0.0093
SH3D21, p = 0.0451
SLAMF7, p = 0.0050
SEC24A, p = 0.0050
SGTB, p = 0.0160
SNAPC1, p = 0.0093
ZC3H12C, p = 0.0025
COPD
control
relative abundance (RT−qPCR)
sputum type
B
orthogonal validation of the most differentially expressed mRNAs in sputum by RT-qPCR
CCL20
R = 0.95 , p = 3.3e−07
CD22
R = 0.93 , p = 2e−06
SH3D21
R = 0.96 , p = 2.2e−07
SLAMF7
R = 0.97 , p = 5.3e−09
SEC24A
R = 0.97 , p = 9.3e−10
SGTB
R = 0.93 , p = 4.7e−07
SNAPC1
R = 0.98 , p = 4.8e−11
ZC3H12C
R = 0.87 , p = 6.3e−05
Sequin normalized read count (mRNA capture sequencing)
relative abundance (RT−qPCR)
sputum type
COPD
control
C
orthogonal validation of the most abundant circRNAs in sputum by RT-qPCR
chr12_69800208_69801721
R = 0.9, p = 2.8e−06
chr14_22909482_22911403
R = 0.88 , p = 6.7e−06
chr1_66891153_66905375
R = 0.98 , p = 8.7e−11
chr2_201145377_201149835
R = 0.94 , p = 6.9e−08
chr3_196391812_196403019
R = 0.76 , p = 0.00062
chr4_143543508_143543977
R = 0.87 , p = 1.4e−05
chr5_137985256_137988315
R = 0.9, p = 2.3e−06
chr6_72295934_72333833
R = − 0.12 , p = 0.67
chr6_72295936_72333835
R = 0.84 , p = 4.6e−05
chr7_103025062_103029418
R = 0.77 , p = 0.0057
Sequin normalized read count (mRNA capture sequencing)
relative abundance (RT−qPCR)
sputum type
COPD
control

**Supplementary Figure 9. Validation of the expression of the most differentially expressed mRNAs and the most abundant circRNAs in sputum by RT-qPCR**

(A) Relative RNA expression of 8 mRNAs based on RT-qPCR in sputum samples of the case- control cohort. Based on RNA sequencing data, all 8 mRNAs are upregulated in sputum from COPD patients compared to sputum from healthy donors. These observations could be validated with RT-qPCR. For each mRNA the p-value for a two-sided Mann-Whitney U test with multiple testing correction using the Benjamini Hochberg method is shown.

(B) Correlation plots of the Sequin normalized read counts and the relative expression based on RT-qPCR for 8 differentially expressed mRNAs in sputum samples of the case-control cohort. The Pearson correlation coefficient and the p-value are shown.

(C) Correlation plots of the Sequin normalized read counts and the relative expression based on RT-qPCR for 10 abundant circRNAs in sputum samples of the case-control cohort. The Pearson correlation coefficient and the p-value are shown.

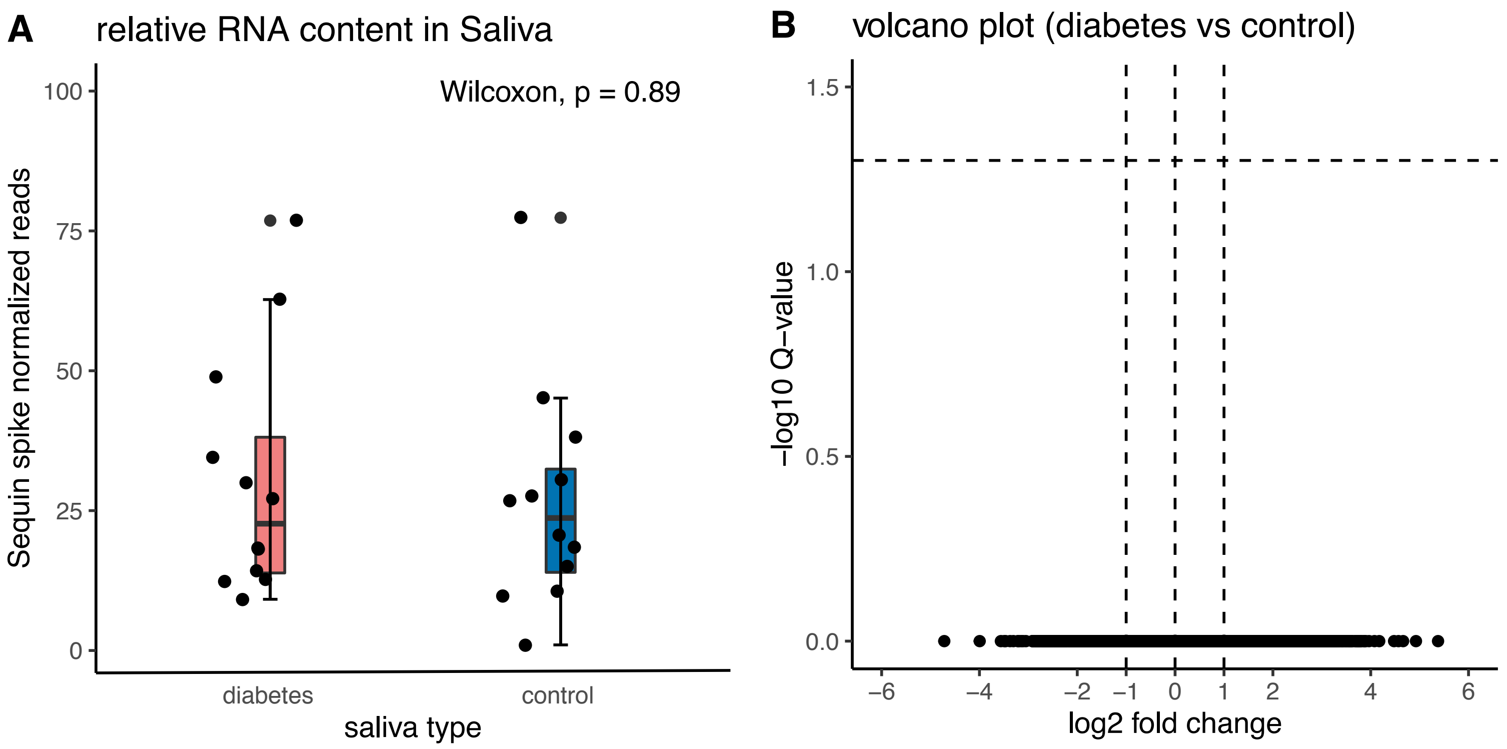

**Supplementary Figure 10. Relative RNA content and volcano plot in the saliva case/control cohort**

(A) The relative RNA content (Sequin normalize reads) is comparable in saliva from diabetes patients (n = 12) and saliva from healthy donors (n = 12; Wilcoxon rank test, two-sided, p = 0.89).

(B) Volcano plot of differentially expressed mRNAs between saliva samples of diabetes patients and saliva samples of healthy volunteers at q < 0.05. No differentially expressed genes were identified.

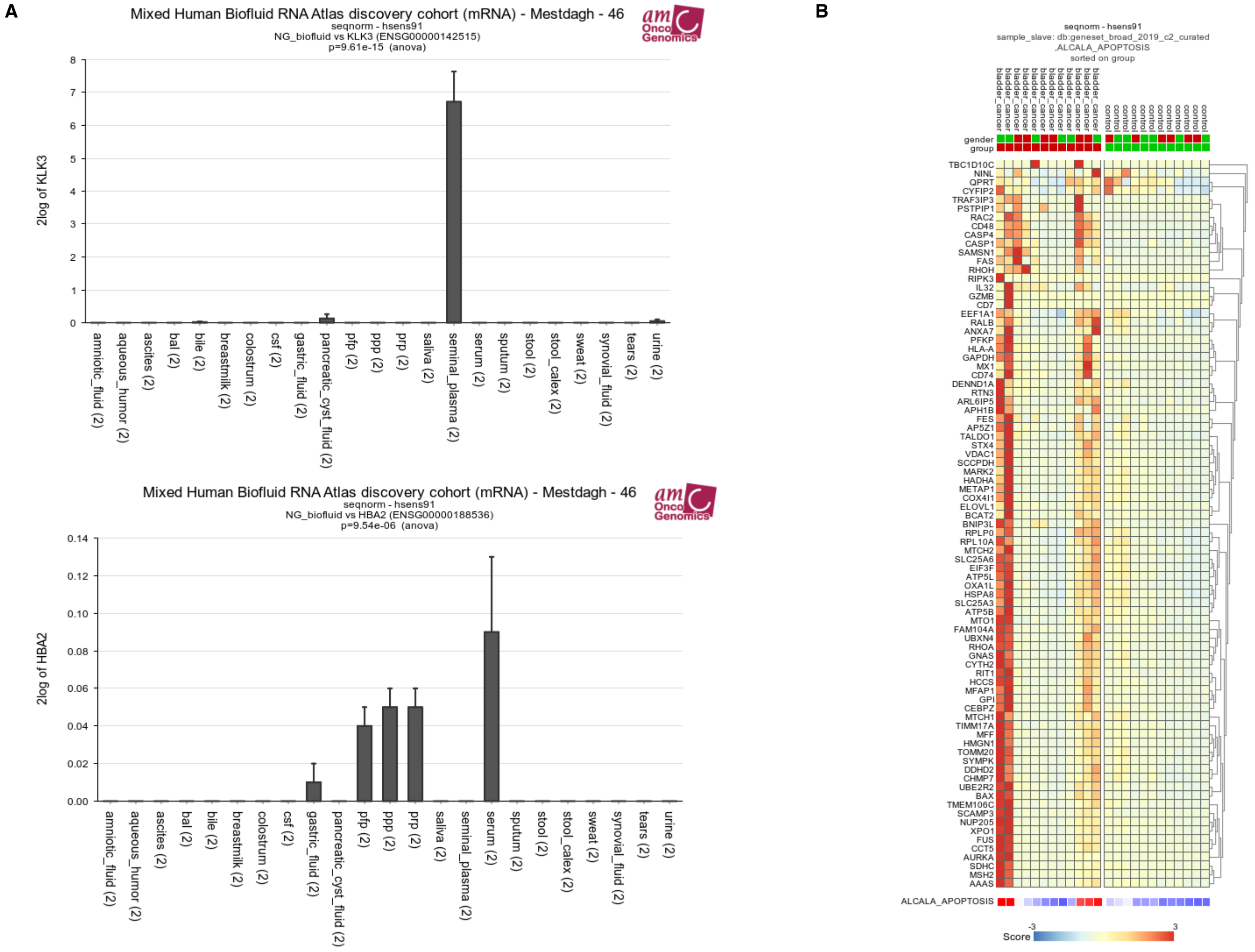

**Supplementary Figure 11. Examples illustrating the applicability of the interactive R2 platform**

Normalized sequencing data can readily be explored in the interactive R2 platform. R2 allows the analysis and visualization of mRNA, circRNA and miRNA abundance. All samples can be used for correlations, principal component analyses, gene set enrichment analysis and many more. Two examples are shown in this panel.

(A) Abundance of KLK3 and HBA2 in the discovery cohort. Prostate-specific antigen is encoded by KLK3. KLK3 is highly abundant in seminal plasma. HBA2 mRNA, coding for the alpha globin chain of hemoglobin, is highly abundant in serum, platelet-rich plasma, platelet-poor plasma and platelet-free plasma.

(B) A heatmap can be generated with a geneset of interest in a specific sample cohort, e.g. the abundance of genes of the ALCALA_APOPTOSIS geneset are plotted per sample of the urine case/control cohort.

### SUPPLEMENTARY NOTES

#### Supplementary Note 1. Details on the RNA spike-in controls

Sequin and external RNA Control Consortium (ERCC) spike-in controls

In the messenger and circular RNA sequencing workflow, Sequin spike-in controls (Garvan Institute of Medical Research, mix A) are added to the biofluid lysate prior to further RNA purification. An overview of the Sequin spikes with their stock concentration is provided in Supplementary Data 1. A volume of 2 µL Sequin spike-in controls at a dilution of 1/1 300 000 was added to 200 µL of all fluid samples, except tears for which an extra pooling step is introduced during the RNA isolation. To enable a fair comparison over all fluids, 2 µL Sequin spike-in controls are added at a dilution of 1/2 600 000 in the tear samples before RNA isolation so that the final pooled RNA samples from tears had the same Sequin amount as the other samples.

ERCC spike-in controls (ThermoFisher Scientific, 4456740, mix 1) are added to the RNA eluate before gDNA removal. In all fluid samples, 2 µL ERCC spike-in controls at a dilution of 1/1 000 000 are added to 12 µL RNA eluate. An overview of the ERCC spikes with their stock concentration is provided in Supplementary Data 2.

RNA extraction Control (RC) and Library Prep control (LP) spike-in controls

In the small RNA sequencing workflow, RNA extraction Control (RC) spike-ins are added to the biofluid lysate prior to RNA isolation, and Library Prep control (LP) spike-in controls to the RNA eluate prior to library prep. All spike-in RNA molecules were ordered at Integrated DNA Technologies as 5'-monophosphate and 3'OH RNA oligonucleotides, and diluted in 500 nM carrier DNA oligo (TCGAAGTATTC). Prior to their use, individual spike-ins are balanced for differences in small RNA sequencing efficiency as described previously(1).

RC spike-in controls are a pool of RNA extraction dynamic range controls (25-mer oligoribonucleotides; RC1-01 - RC1-12; 3-points 10-fold-dilution calibration curve with 3 spikes per dilution) and RNA extraction size controls (25-, 28- or 34-mer oligoribonucleotide; RC2-25 - RC2-34; used at mid-point concentration of the calibration curve) selected from literature(1). RC spike-in IDs and sequences are listed in Supplementary Data 3. A volume of 2 µL from a 190 fM pool (i.e. concentration of highest concentrated spikes that are at 1x in the pool) of RC spikes was added to 200 µL of the lysate before RNA isolation in all fluids, except tears. In tears, 2 µL from a 96.4 fM pool of RC spikes was added before RNA isolation. LP spike-ins are a pool of small RNA sequencing library prep controls (22-mer oligoribonucleotides; LP1-01 - LP1-12; 3-points 10-fold-dilution calibration curve with 3 spikes per dilution) selected from literature.(2) LP spike-in IDs and sequences are listed in Supplementary Data 3. A volume of 2 µL from a 33.8 fM pool (i.e. concentration of highest concentrated spikes that are at 1x in the pool) of LP spikes was added to 12 µL RNA eluate in all fluids.

**Supplementary Note 2. Sample collection protocol per biofluid**

Amniotic fluid

Amniotic fluid was collected in two donors undergoing a clinically-indicated amniocentesis. In one of the donors, the amniocentesis was performed at 22 weeks pregnancy because of CMV seroconversion. No CMV was detected in the amniotic fluid. In the other donor, an amniocentesis was performed at 15 weeks pregnancy because of a *de novo* deletion in 16q22.3 in the firstborn of the parents.

In both donors, 15 mL amniotic fluid was collected in a 50 mL conical centrifuge tube (Falcon, Fisher Scientific). A volume of 1 mL of the amniotic fluid was transferred in a 15 mL conical centrifuge tube. These samples were centrifuged to remove cells (2000 x g, 10 min) and the supernatant was stored at -80 °C within 70 minutes after collection.

Aqueous humor

Aqueous humor was collected in 9 patients undergoing cataract surgery. Topical anesthesia with oxybuprocaine hydrochloride 0.4% was applied to the cornea and conjunctivae. Following instillation of povidone-iodine 5% solution, aqueous humor was obtained through the introduction of a 30-gauge needle on a 1-cc tuberculin syringe onto the corneoscleral junction into the anterior chamber. A volume of 100-200 µL aqueous humor was collected until a shallow anterior chamber was observed. The aqueous humor of donor 1-4 was pooled together obtaining a total volume of 870 µL. The aqueous humor of donor 5-9 was pooled together obtaining a total volume of 850 µL. The two pooled samples were centrifuged to remove cells (2000 x g, 10 min) and the supernatant was stored at -80 °C within 120 minutes after collection.

Ascites (peritoneal fluid) collection

Ascites was collected using paracentesis in two patients with decompensated cirrhosis caused by alcohol (HBV and HCV negative). In each donor, 12 mL ascites was collected with a 20-gauge needle on a 20 mL syringe and transferred to a 15 mL conical centrifuge tube (Fisher Scientific). Both samples were centrifuged to remove cells (2000 x g, 10 min) and the supernatant was stored at -80 °C within 120 minutes after collection.

Bronchoalveolar lavage fluid

Bronchoalveolar lavage fluid (BAL) was collected from two patients during flexible bronchoscopy. In one of the donors bronchoalveolar lavage was performed to obtain microbiological samples for persisting bronchiolitis. In the other donor bronchoalveolar lavage was performed to investigate interstitial lung disease. No infectious focus was found in both samples. Normal saline (100-300 mL divided into three-five aliquots) at room temperature was instilled through the bronchoscope. After the instillation of each aliquot, instilled saline was retrieved using a negative suction pressure less than 100 mm Hg. All aliquots of the retrieved BAL fluid were pooled and 4-5 mfL BAL was transferred to a 15 mL conical centrifuge tube (Fisher Scientific). Both samples were centrifuged to remove cells (2000 x g, 10 min) and the supernatant was stored at -80 °C within 120 minutes after collection.

Bile

Bile was collected in two patients during an endoscopic retrograde cholangiopancreatography (ERCP) procedure. In each donor, 1 mL bile was collected in a sterile recipient before injection of contrast. In the first donor the ERCP was performed for stent exchange in the context of an anastomotic bile duct stricture after liver transplantation. In the second donor, the ERCP was performed for a hilar ischemic cholangiopathy after liver transplantation. All samples were centrifuged to remove cells

(2000 x g, 10 min) and the cell-free supernatant was stored at -80 °C within 2 hours after collection.

Cerebrospinal fluid (discovery cohort)

CSF samples were obtained in two donors to rule out cerebral vasculitis. In both CSF samples normal results were obtained after routine clinical analysis, including the measurement of glucose, total protein content, lactate, lactate dehydrogenase, albumin, IgG, IgM, IgA, leukocytes (with differentiation), erythrocytes and isoelectric focusing. Microbiology cultures were negative in both samples. CSF was aspirated by lumbar puncture using a 25-gauge atraumatic (pencil-point) needle, and transferred to a 15 mL conical centrifuge tube (Fisher Scientific). All samples were centrifuged to remove cells (2000 x g, 10 min) and the cell-free supernatant was stored at -80 °C within 2 hours after collection.

Cerebrospinal fluid (case/control cohort)

Briefly, 4–6 mL of CSF was obtained during a lumbar puncture between the L3 and L5 vertebrae before surgical intervention in glioblastoma patients or during standard therapeutic management of patients with normal-pressure hydrocephalus (non-tumor patients). CSF samples containing blood-derived cells were excluded. Subsequently, CSF samples were centrifuged at 500 g for 10 min at 4 °C, and the cell-free supernatant were aliquoted to 1 mL tubes and stored at −80 °C within one hour after collection.

Gastric fluid

Gastric fluid was collected during oesophagogastroscopy in two patients with complaints of gastroesophageal reflux disease. The procedure was performed after a 12-hour overnight fast. A volume of 4 mL gastric fluid was aspirated using a 10 mL syringe immediately upon entering the stomach. Both samples were centrifuged to remove cells (2000 x g, 10 min) and the cell-free supernatant was stored at -80 °C within 2 hours after collection.

Mature breast milk and colostrum

Colostrum and breast milk were collected in 4 healthy donors using a sterile milk collection unit and an electric breast pump. All samples were transferred to a 15 mL conical centrifuge tube (Fisher Scientific) and centrifuged to remove cells (2000 x g, 10 min). Aliquots of the cell-free supernatant were stored at -80 °C within 50 minutes after collection. Both colostrum samples were collected 3 days after birth. One of the mature breast milk samples was collected 37 days after giving birth, the other sample was collected 50 days after giving birth.

Pancreatic cyst fluid

Pancreatic cyst fluid was collected in two patients during an endoscopic ultrasound procedure. A volume of 1 mL cyst fluid was aspirated with a 10 mL syringe. Both patients underwent the procedure to investigate a cystic lesion in the pancreas. The first patient was diagnosed with a walled off necrosis collection after necrotizing pancreatitis, the second patient was diagnosed with a side-branch intra papillary mucinous neoplasia. The samples were centrifuged to remove cells (2000 x g, 10 min) and stored at -80°C within 2 hours after collection.

Blood draw for plasma and serum

Venous blood was collected in 2 healthy donors from an elbow vein after disinfection with 2% chlorhexidine in 70% alcohol. All blood draws were performed with a butterfly needle of 21 gauge (BD Vacutainer, Push Button Blood Collection Set, Catalogue number 367326, Becton Dickinson and Company, NJ, USA). Tubes were filled to the volume recommended by the manufacturer. Plasma was prepared from whole blood collected in a 10 mL BD Vacutainer K2-EDTA tube (Catalogue number 367525, Becton Dickinson and Company, NJ, USA). Serum was collected in a 6 mL BD Vacutainer SST II Advance tube (Catalogue number

366444, Becton Dickinson and Company, NJ, USA). Immediately after the blood draw, plasma tubes were inverted five times and transported upright in an isolated transportation box at room temperature to the laboratory for further processing. The serum tubes were left on the bench upright for 30-60 minutes to allow clotting, followed by centrifugation at 1200 x g for 15 minutes. The supernatant was subsequently aliquoted per 200 µL into 2 mL LoBind tubes (Eppendorf LoBind microcentrifuge tubes, Z666556-250EA), snap frozen in liquid nitrogen and stored at -80 °C.

Before initiating the first centrifugation step, all plasma tubes were inverted five times. All centrifugation steps were performed in a centrifuge with swinging bucket rotor, without brake and at room temperature.

*Plasma preparation procedure*

Platelet-rich plasma (PRP) was obtained after centrifugation of blood tubes at 400 x g for 20 minutes. Using a 1 mL pipet, plasma was carefully transferred to a fresh 15 mL conical tube (CELLSTAR polypropylene tube, catalogue number 188271, Greiner Bio-One International, Kremsmünster, Austria) leaving approximately 0.5 cm above the buffy coat. If PRP was needed for further experiments, PRP was aliquoted into LoBind tubes (Eppendorf LoBind microcentrifuge tubes, Z666556-250EA), snap frozen in liquid nitrogen and subsequently stored at -80 °C. The remaining volume of PRP was subsequently used for a second spin at 800 x g for 10 minutes. Platelet-poor plasma (PPP) was carefully transferred to a fresh 15 mL tube leaving approximately 0.5 cm above the platelet pellet. If necessary for further experiments, PPP was aliquoted into LoBind tubes (Eppendorf LoBind microcentrifuge tubes, Z666556-250EA), snap frozen in liquid nitrogen and subsequently stored at -80 °C. The remaining volume of PPP was used for a third spin at 2500 x g for 15 minutes. Platelet-free plasma (PFP) was carefully transferred to a fresh 15 mL tube leaving approximately 0.5 cm above the platelet pellet. All PFP was aliquoted in LoBind tubes (Eppendorf LoBind microcentrifuge tubes, Z666556-250EA), snap frozen in liquid nitrogen and stored at -80 °C. All plasma and serum samples were stored at -80 °C within 2 hours after blood collection.

Saliva (discovery cohort)

Spit samples from two healthy donors were collected in a 15 mL conical centrifuge tube. The donors refrained from eating, drinking, smoking and oral hygiene procedures for at least 1 h before sample collection. The samples centrifuged to remove cells (2000 x g, 10 min) and stored at -80 °C within 30 minutes after collection.

Saliva (case/control cohort)

Spit samples from 12 healthy donors and 12 type II diabetes patients were collected in a sterile 15 mL conical centrifuge tube. The donors refrained from eating, drinking other fluids than water, smoking and oral hygiene procedures for at least 1h before sample collection. Five minutes before the collection each donor rinsed his/her mouth twice with water without swallowing it. After a minimum volume of 10 mL saliva per donor was collected, samples were centrifuged (3000 x g, 20 min, 4 °C) and the cell-free supernatant was removed carefully.  Aliquots of 1.5 mL cell-free saliva were stored at -80 °C.

Seminal plasma

Semen samples were produced by masturbation into a sterile container and were allowed to liquefy for 30 min at 37 °C. Samples were centrifuged in a 15 mL conical centrifuge tube to remove cells (2000 x g, 10 min) and stored at -80 °C within 2 hours after collection.

Sputum (discovery cohort)

Spit samples from two healthy donors were collected in a 15 mL conical centrifuge tube. The donors refrained from eating, drinking, smoking and oral hygiene procedures for at least 1h before sample collection. The samples were centrifuged to remove contaminating cells (2000 x g, 10 min) and stored at -80 °C within 30 minutes after collection.

Sputum (case/control cohort)

Sputum induction was performed on 16 subjects who were recruited from the outpatient pulmonary clinic of the Ghent University Hospital or by advertising. Sputum induction and processing was performed as described previously(5). In brief, subjects inhaled sterile, pyrogen-free, hypertonic, nebulised saline at increasing concentrations of NaCl (3%, 4% and 5%) over a 5-minute period after inhalation of salbutamol (2 x 200 µg). Subsequently, subjects were encouraged to cough and expectorate an adequate sample. Sputum plugs were selected, transferred in a polystyrene tube and mixed with dithiotreitol (DTT; 10% Sputalysin with an amount of four times the weight of the sputum plugs, Boehringer-Calbiochem Corp, San Diego, CA, USA) for 30 seconds by vortex and for 15 minutes by tube rocker. Next, an amount of PBS equal to the volume of DTT was added. The sample was incubated for 5 minutes, filtered and centrifuged for separation between cell fraction and cell-free supernatant fraction. The supernatant was aspirated, aliquoted, and stored at -80 °C until further analysis.

Stool

Stool was collected from two healthy donors in a sterile recipient. Two different preparation protocols were followed.

*Protocol without extraction buffer (referred to as "stool" in the manuscript)*

Immediately after collection 0.5 mL stool was transferred from the sterile recipient into a 15 mL conical centrifuge tube. To homogenize the samples 1 mL RNase free Sigma water was added and the samples were vortexed for 30 seconds. The samples were centrifuged to remove cells (2000 x g, 10 min) and stored at -80 °C within 30 minutes after collection.

*Protocol with extraction buffer (referred to as "Calex stool" in the manuscript)*

Immediately after collection stool from the sterile recipient was added to the Calex Cap extraction device (Calex, Bühlmann Laboratories AG, Schönenbuch, Switzerland) according to manufacturer's instructions, resulting in a diluted stool sample (1:500). The diluted sample was subsequently transferred to a 15 mL conical centrifuge tube. The samples were centrifuged to remove cells (2000 x g, 10 min) and stored at -80 °C within 30 minutes after collection.

Sweat

Sweat was collected in two healthy volunteers during exercise on bicycle rollers. Prior to the collection the forehead of the volunteers was disinfected with 2% chlorhexidine in 70% alcohol. Sweat dripping from the forehead was collected in a 15 mL conical centrifuge tube. The samples were centrifuged to remove cells (2000 x g, 10 min) and stored at -80 °C within 30 minutes after collection.

Synovial fluid

Aspiration of synovial fluid from the swollen knee joint was performed in two patients through a sterile puncture with a 21-gauge needle on a 20 mL syringe. The synovial fluid was subsequently transferred to a 15 mL conical centrifuge tube (Fisher Scientific). Both samples were centrifuged to remove cells (2000 x g, 10 min) and stored at -80 °C within 50 minutes after collection. One of the patients was previously diagnosed with spondyloarthropathy, the second patient was diagnosed with rheumatoid arthritis. In both patients, septic arthritis was excluded.

Tear collection

Tears were collected in 8 healthy, non-contact lens wearing volunteers, using Schirmer strips, as previously reported(3,4). Collection was performed under sterile conditions. The closed eyes were first cleaned with a sterile swab. Subsequently 2 Schirmer strips (TearFlo

Sterile Strips, HUB Pharmaceuticals, LLC) per eye were folded into the inferior palpebral fold of the right and the left eye simultaneously. Tears were absorbed into the Schirmer strips while the eyes were closed for maximum 5 minutes. The length of the moistened area was measured using the millimeter scale on the strip. Per donor the total length of the moistened area for ranges from 55 mm to 140 mm (median 127.5 mm). Four Schirmer strips of one donor were collected in one 2 mL Eppendorf tube. RNA isolation was performed within 2 hours after collection.

Urine (discovery cohort)

The mid-stream portion of second morning urine was collected in a sterile recipient. Urine was obtained from one male and one female healthy donor. The samples were centrifuged to remove cells (2000 x g, 10 min) and stored at -80 °C within 30 minutes after collection.

Urine (case/control cohort)

Samples of first morning voided urine were collected in 15 ml tubes with EDTA used for nucleic acid preservation. Samples were centrifuged at 4 °C to remove cells (2000 x g, 15 min). Supernatant was collected and stored at -80 °C until analyzed. Urine was collected in 12 (6 male and 6 female) patients with muscle invasive bladder cancer and in 12 healthy volunteers (6 male and 6 female patients).

### SUPPLEMENTARY TABLES

#### Supplementary Table 1. Demographic and clinical information

Samples were collected in 136 different donors. In general, each sample was obtained from a different donor, with two exceptions:
- sweat, plasma and serum were obtained from 2 healthy volunteers who each provided one sample of the three biofluids
- stool and urine were obtained from 2 healthy volunteers who each provided one sample of both biofluids

| | biofluid | gender (M/F) | mean age (years) | clinical information |
|---|---|---|---|---|
| discovery cohort | amniotic fluid | 0M/2F | 30 | amniocentesis (CMV seroconversion/ de novo deletion in 16q22.3) |
| | aqueous humor | 4M/4F | 72 | cataract surgery |
| | ascites | 1M/1F | 42 | decompensated cirrhosis caused by alcohol |
| | BAL | 1M/1F | 62 | investigation of persisting bronchiolitis/investigation of interstitial lung disease |
| | bile | 1M/1F | 46 | procedure after liver transplantation |
| | CSF | 2M/0F | 51 | lumbal puncture to rule out cerebral vasculitis |
| | gastric fluid | 2M/0F | 51 | gastroesophageal reflux disease |
| | breast milk | 0M/2F | 30 | healthy volunteers |
| | colostrum | 0M/2F | 32 | healthy volunteers |
| | pancreatic cyst fluid | 2M/0F | 68 | investigation of a cystic lesion in the pancreas |
| | plasma and serum | 2M/0F | 34 | healthy volunteers |
| | saliva | 1M/1F | 25 | healthy volunteers |
| | seminal plasma | 2M/0F | 32 | healthy volunteers |
| | sputum | 0M/2F | 30 | healthy volunteers |
| | stool and stool Calex | 1M/1F | 27 | healthy volunteers |
| | sweat | 2M/0F | 40 | healthy volunteers |
| | synovial fluid | 1M/1F | 55 | puncture of the knee joint in patient with spondyloarthropathy/rheumatoid arthritis |
| | tears | 4M/4F | 31 | healthy volunteers |
| | urine | 1M/1F | 27 | healthy volunteers |
| case/control cohort | sputum | 12M/4F | 62 | 8 COPD patients (all smokers) / 8 healthy volunteers (6 smokers and 2 never smokers) |
| | CSF | 12M/12F | 59 | 12 hydrocephalus patients / 12 glioblastoma patients |
| | urine | 12M/12F | 61 | 12 bladder cancer patients / 12 healthy volunteers |
| | saliva | 12M/12F | 48 | 12 diabetes mellitus type II patients / 12 healthy volunteers |

| biofluid case/control cohort | sample ID | Age (y) | gender | ethnicity | group | smoker at moment of collection | years quit smoking | pack year at moment of collection |
|---|---|---|---|---|---|---|---|---|
| sputum | ID94_DMA | 60 | M | Caucasian | control | no | 12 | 48 |
| sputum | RNA006340 | 64 | F | Caucasian | control | no (never smoked) | 0 | 0 |
| sputum | RNA006341 | 61 | M | Caucasian | control | no (never smoked) | 0 | 0 |
| sputum | RNA006342 | 67 | M | Caucasian | control | yes | 0 | 115 |
| sputum | RNA006343 | 52 | F | Caucasian | control | yes | 0 | 56 |
| sputum | RNA006344 | 58 | F | Caucasian | control | yes | 0 | 40 |
| sputum | RNA006345 | 67 | M | Caucasian | control | no | 2 | 57 |
| sputum | RNA006346 | 62 | M | Caucasian | control | no | 13 | 62 |
| sputum | RNA006347 | 71 | M | Caucasian | COPD GOLD II stage | no | 20 | 25 |
| sputum | RNA006348 | 61 | M | Caucasian | COPD GOLD II stage | no | 40 | 1,5 |
| sputum | RNA006349 | 73 | M | Caucasian | COPD GOLD II stage | no | 3 | 19 |
| sputum | RNA006350 | 68 | M | Caucasian | COPD GOLD II stage | no | 15 | 27 |
| sputum | RNA006351 | 64 | M | Caucasian | COPD GOLD II stage | no | 8 | 21,5 |
| sputum | RNA006352 | 46 | M | Caucasian | COPD GOLD II stage | no | 5 | 25 |
| sputum | RNA006353 | 57 | F | Caucasian | COPD GOLD II stage | no | 14 | 20 |
| sputum | RNA006354 | 67 | M | Caucasian | COPD GOLD II stage | no | 8 | 39 |
| CSF | RNA006618 | 64 | M | Caucasian | Glioblastoma | | | |
| CSF | RNA006619 | 43 | M | Caucasian | Glioblastoma | | | |
| CSF | RNA006620 | 47 | M | Caucasian | Glioblastoma | | | |
| CSF | RNA006621 | 59 | M | Caucasian | Glioblastoma | | | |
| CSF | RNA006622 | 41 | M | Caucasian | Glioblastoma | | | |
| CSF | RNA006623 | 64 | M | Caucasian | Glioblastoma | | | |
| CSF | RNA006624 | 51 | F | Caucasian | Glioblastoma | | | |
| CSF | RNA006625 | 69 | F | Caucasian | Glioblastoma | | | |
| CSF | RNA006626 | 59 | F | Caucasian | Glioblastoma | | | |
| CSF | RNA006627 | 68 | F | Caucasian | Glioblastoma | | | |
| CSF | RNA006628 | 57 | F | Caucasian | Glioblastoma | | | |
| CSF | RNA006629 | 76 | F | Caucasian | Glioblastoma | | | |
| CSF | RNA006630 | 66 | M | Caucasian | Hydrocephalus | | | |
| CSF | RNA006631 | 50 | M | Caucasian | Hydrocephalus | | | |
| CSF | RNA006632 | 54 | M | Caucasian | Hydrocephalus | | | |
| CSF | RNA006633 | 72 | M | Caucasian | Hydrocephalus | | | |
| CSF | RNA006634 | 52 | M | Caucasian | Hydrocephalus | | | |
| CSF | RNA006635 | 74 | M | Caucasian | Hydrocephalus | | | |
| CSF | RNA006636 | 57 | F | Caucasian | Hydrocephalus | | | |
| CSF | RNA006637 | 40 | F | Caucasian | Hydrocephalus | | | |
| CSF | RNA006638 | 77 | F | Caucasian | Hydrocephalus | | | |
| CSF | RNA006639 | 56 | F | Caucasian | Hydrocephalus | | | |
| CSF | RNA006640 | 44 | F | Caucasian | Hydrocephalus | | | |
| CSF | RNA006641 | 65 | F | Caucasian | Hydrocephalus | | | |
| urine | RNA006658 | 60 | M | Caucasian | Bladder cancer | | | |
| urine | RNA006659 | 68 | M | Caucasian | Bladder cancer | | | |
| urine | RNA006660 | 72 | M | Caucasian | Bladder cancer | | | |
| urine | RNA006661 | 74 | M | Caucasian | Bladder cancer | | | |
| urine | RNA006662 | 77 | M | Caucasian | Bladder cancer | | | |
| urine | RNA006663 | 84 | M | Caucasian | Bladder cancer | | | |
| urine | RNA006664 | 61 | F | Caucasian | Bladder cancer | | | |
| urine | RNA006665 | 76 | F | Caucasian | Bladder cancer | | | |
| urine | RNA006666 | 52 | F | Caucasian | Bladder cancer | | | |
| urine | RNA006667 | 77 | F | Caucasian | Bladder cancer | | | |
| urine | RNA006668 | 71 | F | Caucasian | Bladder cancer | | | |
| urine | RNA006669 | 53 | F | Caucasian | Bladder cancer | | | |
| urine | RNA006670 | 60 | M | Caucasian | Control | | | |
| urine | RNA006671 | 75 | M | Caucasian | Control | | | |
| urine | RNA006672 | 61 | M | Caucasian | Control | | | |
| urine | RNA006673 | 52 | M | Caucasian | Control | | | |
| urine | RNA006674 | 67 | M | Caucasian | Control | | | |
| urine | RNA006675 | 75 | M | Caucasian | Control | | | |
| urine | RNA006676 | 58 | F | Caucasian | Control | | | |
| urine | RNA006677 | 63 | F | Caucasian | Control | | | |
| urine | RNA006678 | 75 | F | Caucasian | Control | | | |

| | | | | | |
|---|---|---|---|---|---|
| urine | RNA006679 | 62 | F | Caucasian | Control |
| urine | RNA006680 | 67 | F | Caucasian | Control |
| urine | RNA006681 | 49 | F | Caucasian | Control |
| saliva | RNA007039 | 45 | M | Caucasian | diabetes mellitus type II |
| saliva | RNA007040 | 58 | M | Caucasian | diabetes mellitus type II |
| saliva | RNA007041 | 58 | M | Caucasian | diabetes mellitus type II |
| saliva | RNA007042 | 62 | M | Caucasian | diabetes mellitus type II |
| saliva | RNA007043 | 58 | M | Caucasian | diabetes mellitus type II |
| saliva | RNA007044 | 59 | M | Caucasian | diabetes mellitus type II |
| saliva | RNA007045 | 55 | F | Caucasian | diabetes mellitus type II |
| saliva | RNA007046 | 45 | F | Caucasian | diabetes mellitus type II |
| saliva | RNA007047 | 44 | F | Caucasian | diabetes mellitus type II |
| saliva | RNA007048 | 72 | F | Caucasian | diabetes mellitus type II |
| saliva | RNA007049 | 54 | F | Caucasian | diabetes mellitus type II |
| saliva | RNA007050 | 37 | F | Caucasian | diabetes mellitus type II |
| saliva | RNA007051 | 37 | M | Caucasian | control |
| saliva | RNA007052 | 48 | M | Caucasian | control |
| saliva | RNA007053 | 37 | M | Caucasian | control |
| saliva | RNA007054 | 38 | M | Caucasian | control |
| saliva | RNA007055 | 47 | M | Caucasian | control |
| saliva | RNA007056 | 48 | M | Caucasian | control |
| saliva | RNA007057 | 34 | F | Caucasian | control |
| saliva | RNA007058 | 29 | F | Caucasian | control |
| saliva | RNA007059 | 59 | F | Caucasian | control |
| saliva | RNA007060 | 36 | F | Caucasian | control |
| saliva | RNA007061 | 48 | F | Caucasian | control |

**Supplementary Table 2. Tissue samples from the MiOncoCirc database**

| identifier in MiOncoCirc | sample |
|---|---|
| SRR4306580 | bladder cancer 1 |
| SRR4306897 | bladder cancer 2 |
| SRR4307734 | bladder cancer 3 |
| SRR4305726 | bone marrow AML 1 |
| SRR4305798 | bone marrow AML 2 |
| SRR4305921 | bone marrow AML 3 |
| SRR4306287 | bone metastasis 1 |
| SRR4306328 | bone metastasis 2 |
| SRR4307023 | bone metastasis 3 |
| SRR4306139 | cholangiocarcinoma 1 |
| SRR4306159 | cholangiocarcinoma 2 |
| SRR4305774 | colon cancer 1 |
| SRR4306284 | colon cancer 2 |
| SRR4306924 | HNSCC 1 |
| SRR4307034 | HNSCC 2 |
| SRR4306938 | lung cancer 1 |
| SRR4306988 | lung cancer 2 |
| SRR4307026 | lung cancer 3 |
| SRR4307310 | lymph node adenocarcinoma 1 |
| SRR4307322 | lymph node adenocarcinoma 2 |
| SRR1617670 | lymph node breast cancer 1 |
| SRR1617654 | lymph node breast cancer 2 |
| SRR4305892 | medulloblastoma 1 |
| SRR4305911 | medulloblastoma 2 |
| SRR4305928 | medulloblastoma 3 |
| SRR4306449 | melanoma 1 |
| SRR4307654 | melanoma 2 |
| SRR4306559 | melanoma 3 |
| SRR4305758 | neuroblastoma 1 |
| SRR4305938 | ovarian cancer 1 |
| SRR4307799 | prostate cancer 1 |
| SRR4305836 | renal cell carcinoma 1 |
| SRR4306001 | renal cell carcinoma 2 |
| SRR4307062 | urothelial cancer 1 |
