## Supplementary material for "Charting extracellular transcriptomes in The Human Biofluid RNA Atlas": Description of Supplementary Files

**Description of Additional Supplementary Files**

File Name: Supplementary Data 1
Description: Sequin spike-in RNA controls (n=76) during RNA isolation. For each Sequin spike-in, the ID and stock concentration is shown.

File Name: Supplementary Data 2
Description: ERCC spike-in RNA controls during library preparation (n=92). For each ERCC spike-in, the ID and stock concentration is shown.

File Name: Supplementary Data 3
Description: Small RNA extraction Control (RC) spike-ins and Small RNA Library Prep (LP) control spike-ins. For each RC and LP spike-in, the ID, the sequence and the relative concentration is shown.

File Name: Supplementary Data 4
Description: Assessment of tissue and cell contribution to biofluid extracellular RNA is based on publicly available RNA-sequencing data from 27 normal human tissue types and 5 immune cell types (RNA Atlas, Lorenzi et al.). The selection procedure of the tissue types and cell types is explained in "Selection_tissues_celltypes". The gene set per tissue type or cell type is specified in "Markers".

File Name: Supplementary Data 5
Description: Differentially expressed genes in the case/control cohorts using DESeq2 (v1.20.0)

File Name: Supplementary Data 6
Description: Count table with the number of mRNA reads (counts per million) per sample in the discovery cohort

File Name: Supplementary Data 7
Description: Count table with the number of miRNA reads (counts per million) per sample in the discovery cohort

File Name: Supplementary Data 8
Description: Count table with the number of backsplice junction reads (counts per million) per sample in the discovery cohort

File Name: Supplementary Data 9
Description: RT-qPCR assays to validate the expression of the most differentially expressed mRNAs and the most abundant circRNAs in sputum

File Name: Supplementary Data 10
Description: Comparison of the top 10 most abundant miRNAs in amniotic fluid, bronchoalveolar lavage, bile, cerebrospinal fluid, blood plasma, saliva, seminal plasma, serum, urine reported by Godoy et al. with the abundance of these miRNAs in the discovery cohort
